# Unraveling a novel dual-function regulatory element showing epistatic interaction with a variant that escapes genome-wide association studies

**DOI:** 10.1101/2024.03.18.585566

**Authors:** Mathieu Adjemout, Samia Nisar, Amélie Escandell, Hong Thu Nguyen Huu, Magali Torres, Alassane Thiam, Iris Manosalva, Babacar Mbengue, Alioune Dieye, Salvatore Spicuglia, Pascal Rihet, Sandrine Marquet

**Affiliations:** Aix-Marseille Université, INSERM, TAGC, UMR 1090, MarMaRa Institute, Marseille, France; IMI, Institut Pasteur de Dakar, Dakar BP 220, Sénégal; Service D’Immunologie, Université Cheikh Anta Diop de Dakar, Dakar BP5005, Sénégal; Aix Marseille Université, CNRS, Marseille 13009, France; Lead Contact

**Keywords:** Cis-regulatory elements, Enhancer-Silencer, Dual function, Epistatic interaction, Functional variants, Severe malaria, T cells activation, Transcriptional regulation

## Abstract

Regulation of gene expression has recently been complexified by the identification of Epromoters, a subset of promoters with enhancer function. Here, we uncovered the first dual cis-regulatory element, "ESpromoter," exhibiting both enhancer and silencer function, as a regulator of the nearby genes *ATP2B4* and *LAX1* in single human T cells. Through integrative approach, we pinpointed functional rs11240391, a severe malaria risk variant that escapes detection in genome-wide association studies, challenging conventional strategies for identifying causal variants. CRISPR-modified cells demonstrated the regulatory effect of ESpromoter and rs11240391 on *LAX1* expression and T cell activation. Furthermore, our findings revealed an epistatic interaction between ESpromoter SNPs and rs11240391, impacting severe malaria susceptibility by further reducing *LAX1* expression. This groundbreaking discovery challenges the conventional enhancer-silencer dichotomy. It highlights the sophistication of transcriptional regulation and argues for an integrated approach combining genetics, epigenetics, and genomics to identify new therapeutic targets for complex diseases.

**HIGHLIGHTS:**

- Novel dual enhancer-silencer element (ESpromoter) in a single human cell type
- Functional SNP for severe malaria risk that escapes genome-wide association studies
- Genome editing at the SNP demonstrates a regulatory effect on *LAX1* and T cell activation
- Epistatic interaction between SNPs increases the risk of severe malaria

**In brief:** Epistatic interaction between common variants within a novel dual enhancer-silencer regulatory element and the *LAX1* promoter variant is responsible for severe malaria susceptibility through T-cell activation.

## INTRODUCTION

Gene expression is tightly regulated in time and space to ensure that genes are switched on or off at the right times and in the right amounts. Disruption in this regulation can result in a wide range of diseases and syndromes including cancers^1,2^, neurological^3,4^, developmental disorders, autoimmune, cardiovascular, and infectious diseases^5^. Gene expression profiling has therefore emerged as a powerful approach for gaining deeper insights into the pathological mechanisms of complex diseases by identifying specific transcriptomic signatures linked to these pathologies^6–13^.

In humans, the regulation of gene expression occurs through the intricate interplay between proximal promoters located near the 5’ end of genes, responsible for initiating their expression, and distal cis-regulatory elements (CREs), known as enhancers^14^. These enhancers can activate gene transcription over large distances and independently of orientation. Recent studies have unveiled that a subset of promoters can also serve as enhancers, termed “Epromoters”^15–18^. Investigating these promoters, enhancers or Epromoters is essential for understanding complex gene regulation, as they wield the power to directly influence physiological traits or diseases by simultaneously regulating the expression of numerous proximal and distal genes. Furthermore, the presence of sequence variations within these regulatory elements adds another layer of molecular complexity. These variations have the potential to directly impact disease development by altering the binding sites for transcription factors, leading to the deregulation of target gene expression. The cis effects of non-coding variants on gene expression could therefore be the main risk factor for complex phenotypic traits and disease susceptibility as supported by genome-wide association studies (GWAS). Over 90% of the variants uncovered through GWAS are located in non-coding regions of the genome, frequently lacking comprehensive functional annotations^19,20^. Consequently, the study of the functional significance of specific risk variants is becoming essential to elucidate the underlying pathogenesis in the fields of biology and health. Despite the identification of numerous genomic loci associated with severe malaria, the precise causal genetic variants, the genes responsible and the mechanisms by which they operate remain largely unknown. Identifying these regulatory variants and elucidating their functional implications is therefore a major challenge requiring new approaches.

To address this challenge, we have devised an innovative strategy aimed at identifying within regulatory elements, causal variants associated with severe malaria by dissecting their impact on gene expression. This study is of paramount importance in improving our understanding of the regulatory mechanisms governing malaria susceptibility genes, given the persistent threat to public health posed by malaria. This knowledge is essential for the development of innovative, effective, and sustainable therapeutic interventions. In this context, we have established a comprehensive pipeline that integrates genetic and epigenomic data using a combination of bioinformatics and experimental approaches. Our previous research resulted in the identification of an Epromoter within the *ATP2B4* gene locus^17^, demonstrating the efficacy of our strategy. This Epromoter comprises five regulatory variants -namely rs11240734, rs1541252, rs1541253, rs1541254, and rs1541255-all of which have been associated with severe malaria^17,21^ and are in strong linkage disequilibrium (LD) with the non-functional lead SNP (rs10900585) previously identified by genome-wide association studies (GWAS)^22–26^. The current challenge is to establish the link between this disease-associated region and the different genes affected, while also identifying the specific cell types involved. This is essential for unraveling the mechanisms and biological pathways that underlie the genetic susceptibility of individuals to malaria.

Here, our aim was therefore to assess the ability of this *ATP2B4* Epromoter to regulate the expression of other genes within this same genomic region, potentially involved in malaria susceptibility. Thanks to this study, we have discovered the first dual cis-regulatory element exhibiting both an enhancer (E) activity on the *ATP2B4* gene and a silencer (S) activity on the *LAX1* gene in the same human cell type, we called “ESpromoter”.

Subsequently, we have demonstrated that the silencing function of this ESpromoter on the *LAX1* target gene is influenced by the presence of a functional variant (rs11240391) located in its promoter region, revealing an epistatic interaction between them. This causal variant, which is not in LD with any other polymorphism in African populations, cannot be detected through Genome-Wide Association Studies (GWAS) due to its absence among genotyped Single Nucleotide Polymorphisms (SNPs). Furthermore, our research also highlights the limitations inherent in the exclusive use of GWAS strategies for the identification of causal variants. We illustrate that our more comprehensive approach, integrating multi-omics data, proves more effective in identifying regulatory elements and variants.

## RESULTS

### *LAX1* is a target of *ATP2B4* Epromoter

Characterized as Epromoter, the *ATP2B4* internal promoter can potentially interact with other genes, even those hundreds of kilobases away. Prediction of the interaction between Genehancer regulatory elements and neighboring genes suggested that the *ATP2B4* Epromoter may have four target promoters, including *ATP2B4*, *LAX1*, *ZC3H11A,* and *OPTC* (Figure 1A). Promoter capture Hi-C data from CD4+ T cells^27^, revealed a significant T-cell specific interaction between *ATP2B4* Epromoter and both *LAX1* and *ATP2B4* promoters (Figure 1B, Figure S1). However, no significant interaction was detected between the *ATP2B4* Epromoter and the *ZC3H11A* and *OPTC* genes in CD4+ T cells^27^.

**Figure 1.**
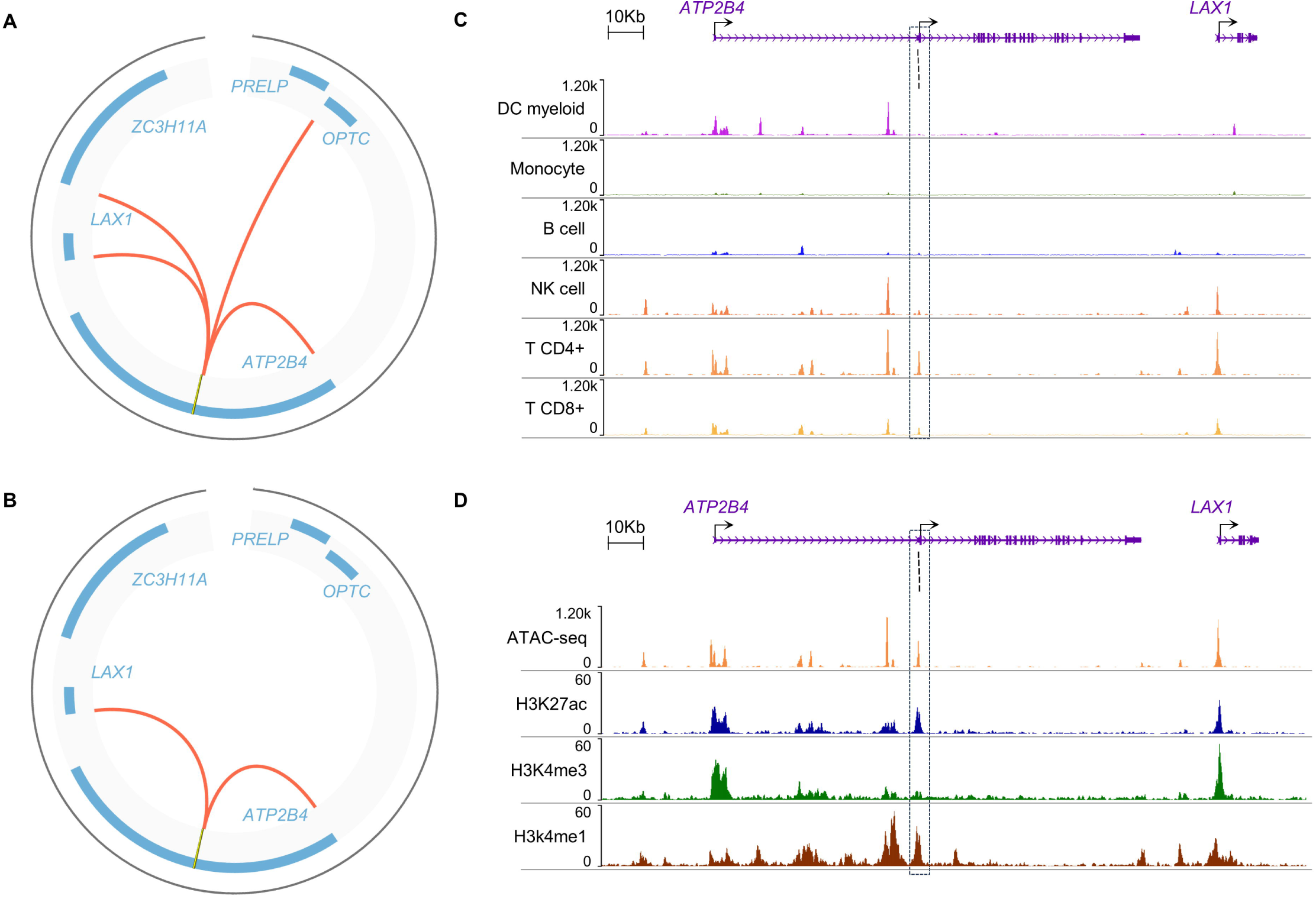
Epromoter in the *ATP2B4* locus, a potential cis-regulatory element of *LAX1*. (A) Circos plots showing predictive chromatin interactions of Genehancer regulatory elements with the *ATP2B4* Epromoter (yellow line). (B) Circos plots showing significant chromatin interactions with the *ATP2B4* Epromoter (yellow line) in primary T CD4+ cells as measured by promoter capture Hi-C. (C) WashU epigenome browser view of ATAC-seq for immune cells. The frame corresponds to the Epromoter. Black lines correspond to the 5 SNPs previously identified (rs11240734, rs1541252, rs1541253, rs1541254, and rs1541255) (D) WashU epigenome browser view of epigenomic data in naive CD4+ primary cells. The frame corresponds to the Epromoter. Black lines correspond to the 5 SNPs previously identified (rs11240734, rs1541252, rs1541253, rs1541254, and rs1541255)

### *ATP2B4* Epromoter and the *LAX1* promoter are active chromatin regions in CD4+ T cells

To deeper characterize the epigenetic context of *ATP2B4* in immune cells, we looked at chromatin features that may be particularly useful for identifying potential regulatory regions and their cell-specific function. More, specifically, the study of chromatin accessibility using ATAC-seq (Assay for Transposase-Accessible Chromatin sequencing) profiles in various human immune cells^28^ confirmed the existence of cell-specific open chromatin in the *ATP2B4* locus (Figure 1C). Particularly high accessibility was observed in CD4+ T cells for the cis-regulatory region containing malaria-associated variants and for the *LAX1* promoter. These two regions also showed chromatin accessibility in CD8+ T cells and Natural Killer (NK) cells (Figure 1C). These open regions could serve as binding sites for transcription factors, facilitating cell-specific regulation. We have also studied other epigenetic features such as histone marks, in CD4+ T cells, where an interaction between these two regions was established. The analysis of acetylation profiles of histone 3 lysine 27 (H3K27Ac) profiles from Roadmap data^29^, accessible via the WashU genome browser, revealed active gene regulatory elements corresponding to the *ATP2B4* Epromoter and the *LAX1* promoter in CD4+ T cell (Figure 1D). Notably, malaria-associated SNPs were found to align with an H3K27Ac peak in CD4+ T cells. In addition, the *ATP2B4* Epromoter showed H3K4me1 mark stronger than H3K4me3 mark, whereas the *LAX1* promoter exhibited a more pronounced H3K4me3 mark. These observations are consistent with their respective roles as enhancer or promoter in this specific cell type. These characterizations confirm that the *ATP2B4* Epromoter and the *LAX1* promoter are active chromatin regions (ACRs) in CD4+ T cells. Overall, the data indicate a strong correlation between DNA accessibility and histone marks, supporting T cells as a pertinent model for investigating the underlying regulatory mechanisms. Interestingly, similar profiles were observed for Jurkat cells, a human T cell line (Figure S2), which prompted us to use them for further experiments aimed at elucidating the regulatory mechanisms governing these regions.

### The *ATP2B4* cis-regulatory element has a dual enhancer-silencer function in a single cell type

To determine the regulatory function of the *ATP2B4* CRE on the target genes (*ATP2B4* and *LAX1*), we performed genome editing experiments in the Jurkat cell line. We deleted a 506 bp region encompassing the rs11240734, rs1541252, rs1541253, rs1541254, and rs1541255 SNPs using CRISPR/Cas9 with two flanking guide RNAs (gRNAs) (Figure 2A). Three clones with a 506 bp biallelic deletion (Δ1, Δ2, and Δ3) were generated from a total of 457 screened clones. An unedited wild-type clone, also exposed to the CRISPR/Cas9 complex was selected. Expression of these genes was assessed without and with stimulation, since *LAX1* which is involved in TCR signaling requires PMA/ionomycin stimulation to be induced. In the wild-type, a slight decrease in the expression of long *ATP2B4* transcripts was observed under PMA/ionomycin stimulation (Figure 2B) as previously observed^30^. Deletion of the 506 bp region resulted in decreased expression of the two long *ATP2B4* transcripts corresponding to ENST00000367218 and ENST00000357681, in the presence or absence of PMA/ionomycin stimulation (Figure 2B), with a 1.8-, 1.2- and 2.9-fold decrease for Δ1, Δ2, and Δ3 respectively in stimulated condition (p < 0.01).

**Figure 2.**
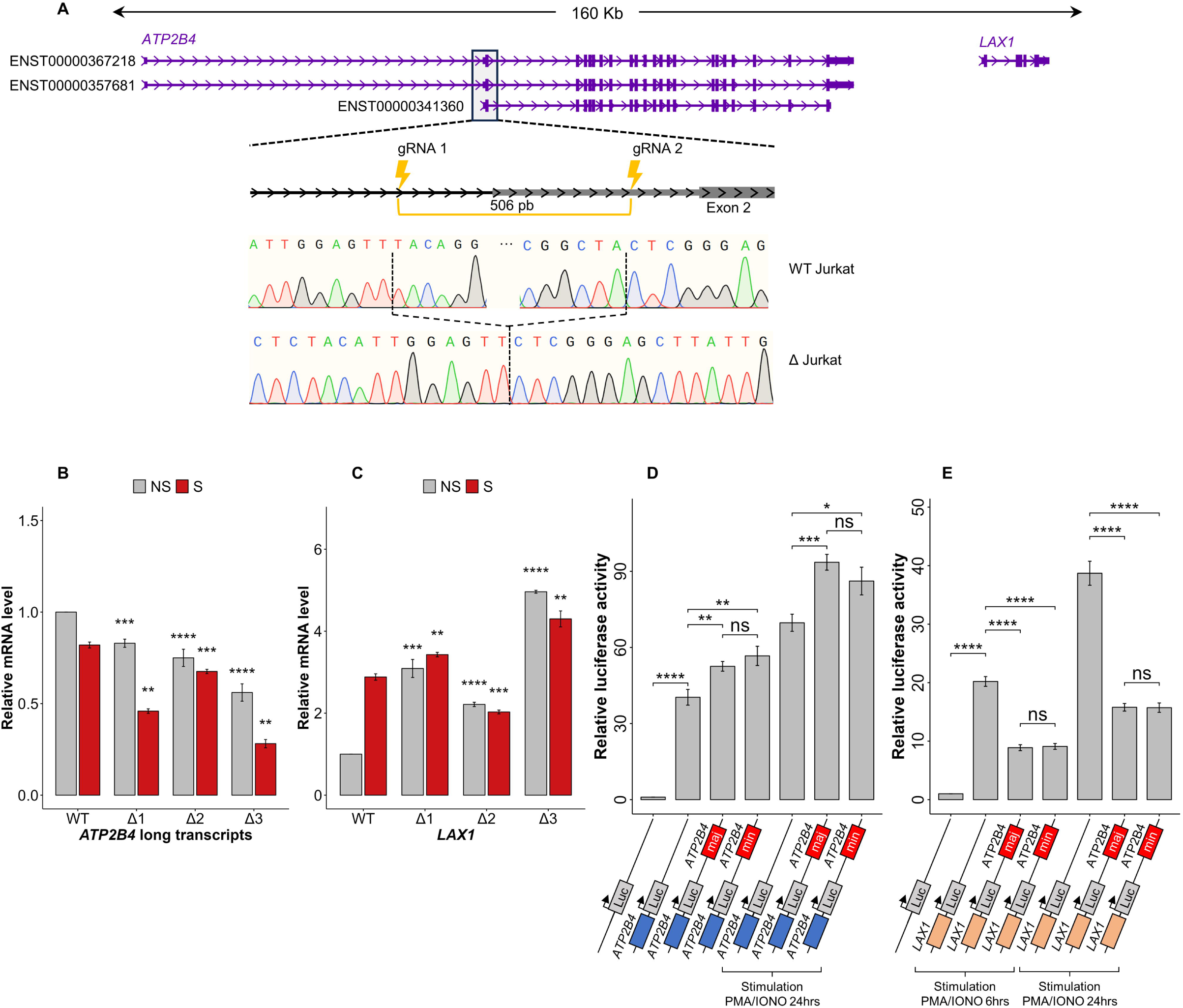
Identification of a new cis-regulatory element with a dual enhancer-silencer function (ESpromoter) (A) Generation of cell lines with a 506-bp deletion of the ESpromoter using two single guide RNAs (gRNA1 and gRNA2) flanking the regulatory region containing the 5 SNPs through CRISPR-Cas9 technology. The position of the deletion is indicated relative to the long *ATP2B4* (ENST00000367218, ENST00000367218) and short (ENST00000341360) transcripts of *ATP2B4* and the *LAX1* gene. Sanger sequencing chromatograms show the genomic sequence of the wild-type Jurkat clone (WT Jurkat) and the edited clone (Δ Jurkat). (B) qPCR analysis of *ATP2B4* long transcript expression (ENST00000357681, ENST00000367218) on wild-type Jurkat cells and clones deleted for the ESpromoter (Δ1, Δ2, and Δ3) in culture without stimulation (NS: no stimulation in grey) or with PMA/ionomycin stimulation for 6 hours (S: stimulated in red). Clones with a deletion had a decreased *ATP2B4* expression under both conditions. Values were generated from 3 independent experiments. Wild-type Jurkat ns and s are used as references. (C) qPCR analysis of *LAX1* expression on wild-type Jurkat cells and on clones deleted for the ESpromoter (Δ1, Δ2, and Δ3) without stimulation (NS: no stimulation in grey) or with PMA/ionomycin stimulation for 6 hours (S: stimulated in red). Values were generated from 3 independent experiments. Clones with a deletion had increased *LAX1* expression under both conditions. Wild-type Jurkat NS and S are used as references. (D) Graphs showing the relative luciferase activity under the control of *ATP2B4* promoter alone or in combination with the ESpromoter region containing either the major haplotype (maj: TCCGA) or minor haplotype (min: CTTGG) for the 5 SNPs (rs11240734, rs1541252, rs1541253, rs1541254, and rs1541255) without and with PMA/ionomycin stimulation. Luciferase assays confirmed the enhancer effect of the ESpromoter on the *ATP2B4* promoter independently of the haplotype. (E) Graphs showing the relative luciferase activity under the control of *LAX1* promoter alone or in combination with the ESpromoter region containing either the major haplotype (maj) or minor haplotype (min) for the 5 SNPs (rs11240734, rs1541252, rs1541253, rs1541254, and rs1541255) under PMA/ionomycin stimulation (6 h and 24 h). Luciferase assays confirmed the silencer effect of the ESpromoter on the *LAX1* promoter independently of the haplotype. All data represent mean values ± SEM of three independent experiments. P-values were calculated using a two-sided Student’s t-test.

As shown in Figure 2C, the *LAX1* gene has a PMA/ionomycin-inducible promoter whose expression increased 3-fold in wild-type Jurkat cells after stimulation. However, *LAX1* gene expression increased significantly in all three clones with the 506 bp deletion, in the presence or absence of stimulation. Thus, the *LAX1* gene is highly expressed in Δ1, Δ2 and Δ3 clones, even in the absence of stimulation (Figure 2C).

These data demonstrate that deletion of the *ATP2B4* cis-regulatory element simultaneously results in decreased of *ATP2B4* expression and increased expression of *LAX1* in the Jurkat cell line. These results suggest that the cis-regulatory region acts as an enhancer for *ATP2B4* and as a silencer for *LAX1*. We have therefore identified a new type of regulatory element that we named “ESpromoter”, which has the dual function of both enhancer and silencer in the same cell type (Figure S1).

### The dual enhancer-silencer function is independent of the variants within the ESpromoter

To confirm the dual function of the cis-regulatory element of *ATP2B4* (ESpromoter) and assess the impact of SNPs on gene regulation, we performed luciferase gene reporter assays in the Jurkat cell line. First, we transfected the cells with a construct containing the ESpromoter region downstream of the luciferase reporter gene, and the 780-bp *ATP2B4* long transcript promoter upstream of luciferase (Figure 2D). We initially validated the promoter activity of this 780-bp region compared to the basic vector (p < 0.0001, with a 40-fold increase). We also confirmed the enhancer activity of the 601-bp ESpromoter containing the five SNPs, on the *ATP2B4* promoter without stimulation (p < 0.01, with a 1.3-fold increase) and with PMA/ionomycin stimulation for 24 hours. However, no significant difference was observed between the constructs with minor or major alleles for the 5 SNPs located in the ESpromoter (Figure 2D). We then transfected Jurkat cells with constructs in which the *ATP2B4* promoter was replaced by the 810-bp *LAX1* promoter region. We showed the activity of the *LAX1* promoter after 6 h or 24 h of PMA/ionomycin stimulation compared to the basic vector (p < 0.0001, fold increase of 20 for 6 hours and 39 for 24 hours) (Figure 2E) as well as the silencing effect of the ESpromoter region on *LAX1* promoter (p < 0.0001, fold decrease of 2.24 for 6 hours and 2.45 for 24 hours). The decrease in expression was the same for both major and minor haplotypes. These data confirm that this regulatory region exhibits enhancer and silencer activity independent of *ATP2B4* SNP alleles in the Jurkat cell line. We have therefore confirmed the existence of a new type of regulatory element “ESpromoter” with a dual enhancer-silencer function.

### Increased expression of *LAX1* in clones lacking the ESpromoter region delayed T-cell activation

Since the adaptor protein LAX functions as a negative regulator of lymphocyte signaling^31^, an increase in *LAX1* expression is expected to prevent T lymphocyte activation. To evaluate the activation of Jurkat wild-type cells and Jurkat clones lacking the ESpromoter region after PMA/ionomycin stimulation, we followed the surface expression of CD69, a T cell surface activation marker rapidly induced after stimulation through the T cell antigen receptor (TCR) as a measure of T cell activation. Jurkat unmodified cells and deleted clones were stimulated for 4 hours with PMA/ionomycin, and CD69 cell surface expression was measured before stimulation, at 15-minute intervals between 1 and 2 hours, and at the end of the 4-hour period after stimulation (Figure 3A, 3B, 3C). We showed that the number of CD69-positive cells was similar for wild-type Jurkat cells, and the unedited wild-type clone exposed to CRISPR/Cas9 (WTc) at all stimulation times (Figure 3A and 3B). The number of CD69-positive cells increased with stimulation time for the WT, WTc, and Δ3. However, at each time point, the number of CD69-positive cells was higher for unmodified cells (WT and WTc) compared to clone Δ3, with around 90% and 70% respectively after 4 h of stimulation. This difference was statistically significant using a paired t-test (p = 0.001) (Figure 3A and 3C).

**Figure 3.**
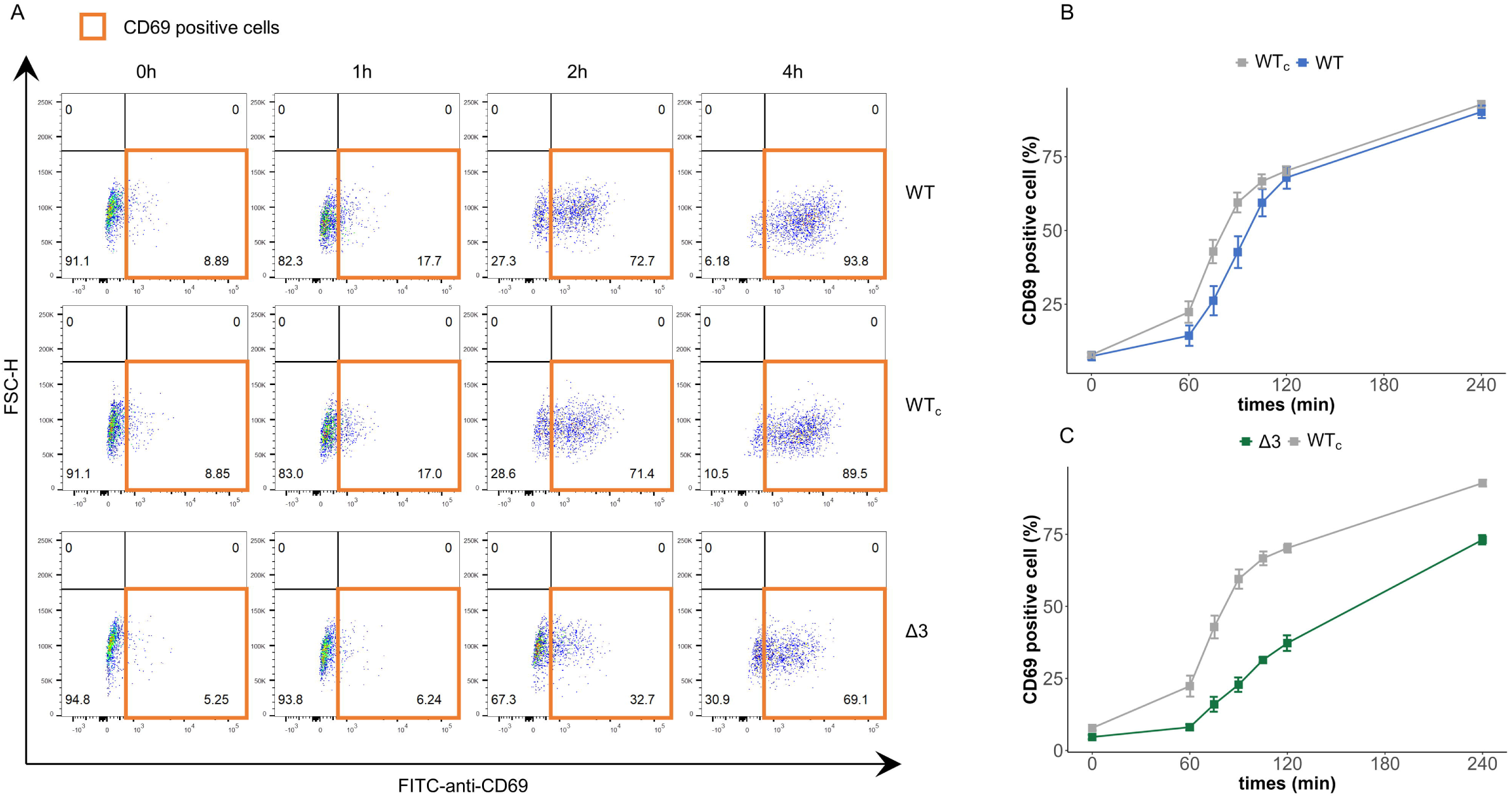
Inhibition of T-cell activation in the ESpromoter-deleted clone. (A) T cell activation was assessed based on CD69 staining and flow cytometric gating strategy. Plots showing the results at different time points after PMA/Ionomycin stimulation of wild-type Jurkat cells (WT), wild-type clone after Crispr/Cas9 editing (WTc), and deleted clone for the ESpromoter (Δ3). Representative experiments on 2000 of each type in function to Forward Scatter-Horizontal (FSC-H) and anti-CD69 staining with Fluorescein Isothiocyanate (FITC). (B) Monitoring of Jurkat cell activation by anti-CD69 staining through flow cytometry after PMA/ionomycin stimulation. Values represent the average ± SEM of two independent experiments performed in duplicate, indicating the percentage of cells positive for anti-CD69 staining (acquisition of 2000 cells per clone) for wild-type Jurkat (WT), and wild-type clone without genomic edition after CRISPR-Cas9 (WTc). A similar percentage of CD69-positive cells was observed between WT and WTc. (C) Monitoring of Jurkat cell activation by anti-CD69 staining through flow cytometry after PMA/ionomycin stimulation. Values represent the average ± SEM of two independent experiments performed in duplicate, indicating the percentage of cells positive for anti-CD69 staining (acquisition of 2000 cells per clone) for wild-type clone without genomic edition after CRISPR-Cas9 (WTc) and deleted clones for the ESpromoter (Δ3). A lower number of CD69-positive cells was observed in the Δ3 clone.

### Discovery of the functional variant rs11240391 in the *LAX1* promoter

Non-coding variants can affect gene expression by modifying enhancer function, thereby regulating cell-specific gene expression. However, here we found no evidence of allele-specific regulation of *LAX1* by the five *ATP2B4* SNPs located in the ESpromoter. We therefore investigated whether SNPs within the *LAX1* promoter, identified as the target region of the ESpromoter through Hi-C experiments (Figure 1B), could indeed functionally regulate its expression. To identify such regulatory SNPs, we used various databases and tools to prioritize the most promising candidates. Hence, we first searched for expression Quantitative Trait Loci (eQTLs) using the Elixir eQTL catalogue^32^, revealing several SNPs with a PIP (Posterior Inclusion Probability) score > 0.2 (Figure 4A), indicating the probability of the variant being functional. Among them, rs11240391, located in the *LAX1* promoter, exhibited a high PIP score of 0.76 (p = 1.09 x 10^-10^) in whole blood tissue (Figure 4A). This SNP is also described as an eQTL for *LAX1* in CD4+ T cells, with a PIP of 0.92, in FIVEx database^33^. Furthermore, the ENCODE data confirm that the *LAX1* promoter region corresponds to a cis-regulatory element (Figure 4A), where numerous Chip-seq peaks for transcription factors were identified using ReMap2022^34^ (Figure 4A). We therefore focused on further experiments to elucidate the cis-regulation role of rs11240391 in Jurkat cells. To study its impact on *LAX1* expression, we cloned the *LAX1* promoter with each allele into a luciferase reporter construct and measured relative expression in the presence or absence of PMA/ionomycin stimulation. Upon stimulation, luciferase expression increased for both the T and G alleles (Figure 4B).

**Figure 4.**
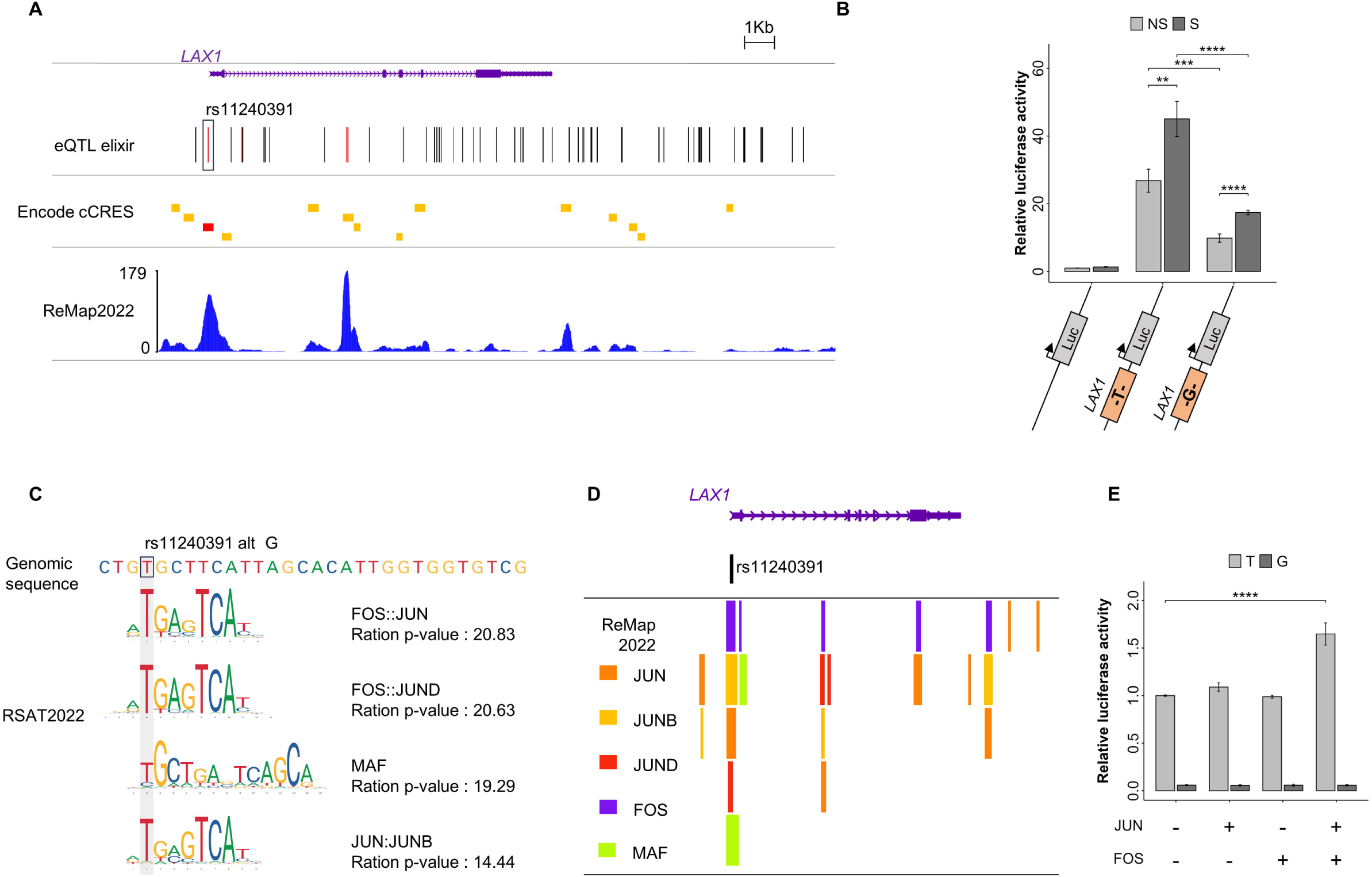
Identification of the functional variant rs11240391 corresponding to the FOS and JUN binding site. (A) WashU epigenome browser view of *LAX1* region with the position of all eQTLs for *LAX1* in the ELIXIR database, those having a PIP score > 0.7 are shown in red. Localization of Encode cCRES was shown with promoter-like elements in red and enhancer-like elements in orange. The density of CHIP-seq peaks available in ReMap2022 is also displayed. Overall, these results revealed a potentially functional SNP in the *LAX1* promoter (rs11240391). (B) Luciferase assays to assess the impact of the rs11240391 variant on *LAX1* promoter activity without stimulation (NS: no stimulation in grey) or with PMA/ionomycin stimulation for 6 hours (S: stimulated in black). Relative luciferase activity was lower with the “G” allele than with the “T” allele. The graph shows the mean values ± SEM. P-values were calculated using a two-sided Student’s t-test. (C) Prediction of transcription factor binding sites disruption at rs11240391 using RSAT. The genomic region containing the variant is displayed. The P-value ratio was calculated by dividing the best probability of transcription factor binding by the worst probability. All transcription factors exhibit a higher binding affinity to the T allele of SNP rs11240391. (D) CHIP-seq peaks from ReMap2022 confirmed the binding of transcription factors, identified by RSAT, in the region containing rs11240391. (E) Luciferase assays to evaluate the impact of rs11240391 on the fixation of transcription factors by adding expression plasmid of JUN or FOS or both. The major allele T was represented in grey, and the minor allele G was represented in black. A higher relative luciferase activity was observed in the presence of the "T" allele, which only increased with the simultaneous addition of JUN and FOS. Values were generated in triplicate from 3 independent experiments. The plot shows the mean values ± SEM. P-values were calculated using a two-sided Student’s t-test.

However, the minor “G” allele at rs11240391 showed reduced expression compared to the major “T” allele under both conditions (p < 0.001) (Figure 4B) confirming the functional role of this SNP (Figure S1).

### The regulatory function of the rs11240391 SNP was mediated by the FOS and JUN transcription factors

Transcription factors (TFs) are known to be key players in gene regulation, and their binding to target regulatory regions can be altered by genetic variants. To uncover the regulatory mechanism and determine whether TF binding may be different between rs11240391 alleles, we used the RSAT tool. Hence, we identified *in silico* several factors whose binding appeared to be affected by this SNP (Figure 4C), suggesting preferential binding of these factors on the “T” allele. The best p-value ratio was obtained for FOS:JUN (p = 20.83). The use of ChIP-seq data from ReMap 2022^34^ showed the binding of the TFs JUN, JUNB, JUND, FOS, and MAF to the *LAX1* promoter (Figure 4D). These data suggest a potential role for FOS and JUN in the regulation of the *LAX1* gene. Therefore, we next focused on these two TFs and assessed whether the rs11240391 variant of the *LAX1* promoter could be a potential target of FOS and JUN. Biological relevance was investigated by luciferase assays in Jurkat cells. As shown in Figure 4E, in the presence of plasmids expressing the FOS and JUN proteins, relative luciferase activity was significantly increased (p < 0.0001, 1.69-fold increase) only in the presence of the T allele of the rs11240391 variant. This result suggests that the FOS and JUN transcription factors can bind to the *LAX1* promoter specifically on the T allele as predicted by RSAT (Figure 4C). Thus, a clear potentialization of the activation effects can be observed in the presence of FOS and JUN allowing the formation of the AP1 dimer.

### Genetic editing of the rs11240391 allele confirms its regulatory impact on *LAX1* expression

To confirm whether the effect on *LAX1* gene expression is allele-specific at rs11240391, we performed homologous recombination (HR) with CRISPR/Cas9 genome editing to replace the T alleles of wild-type Jurkat cells by the G allele (Figure 5A). After screening more than 700 clones, only two heterozygous (T/G) clones were successfully recovered through HR without any additional insertions, deletions, or base modifications (Figure 5A). The limited efficiency of HR with CRISPR/Cas9 is likely attributed to the difficulty of deactivating the guide RNA (gRNA) once HR has been accomplished. Notably, the Cas9 enzyme can cleave the integrated DNA fragment again, as the protospacer adjacent motif (PAM) remains unchanged. This precaution is taken to avoid introducing a new mutation by altering the PAM sequence on the DNA donor carrying the minor allele T. *LAX1* quantification was performed by RT-qPCR, showing a significant decrease of the *LAX1* transcript in both heterozygous clones compared with the wild-type cells unmodified after CRISPR/Cas9 (WT_HR_) (p < 0.001, 2.12-fold decrease) (Figure 5B).

**Figure 5.**
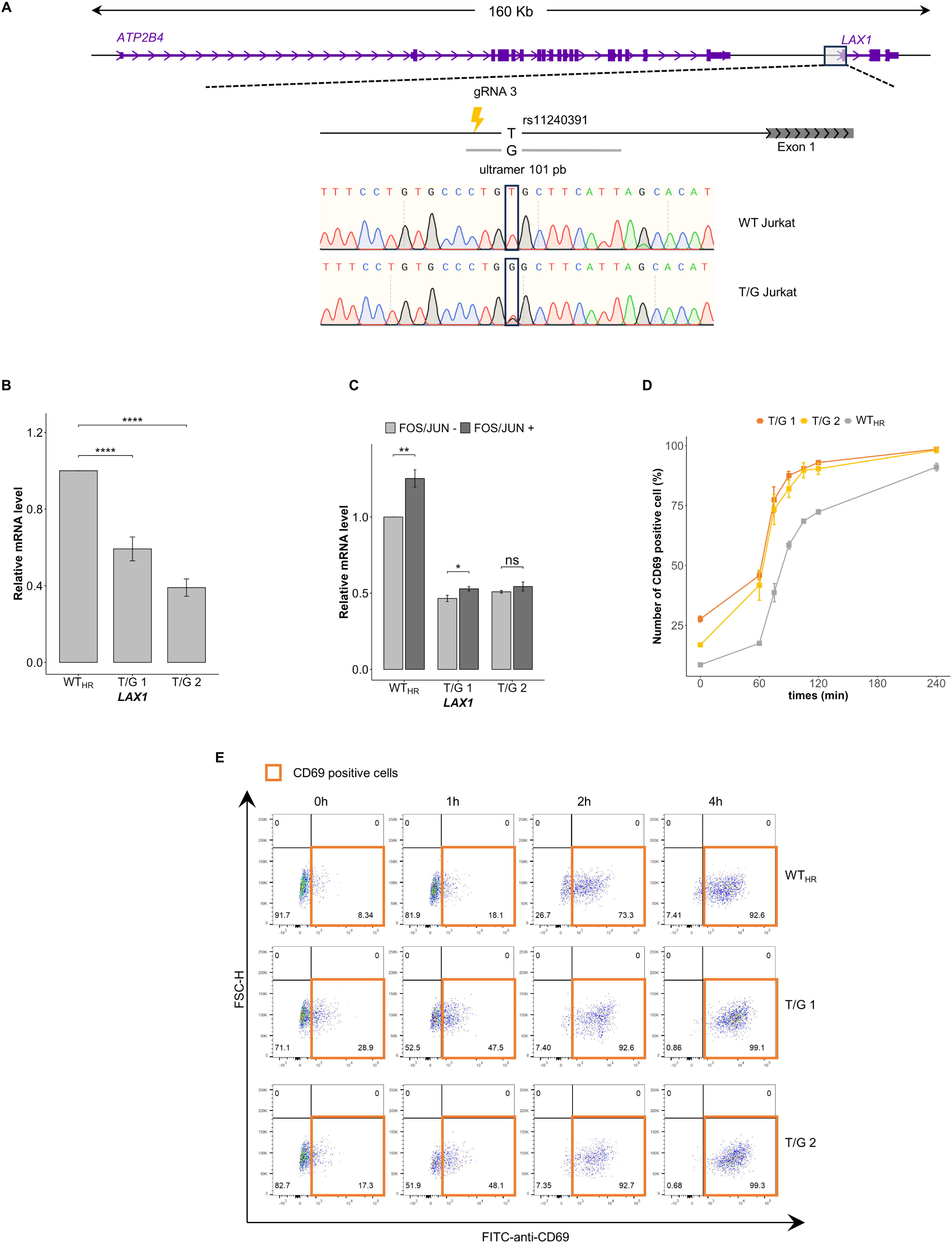
Lower *LAX1* expression in heterozygous T/G clones is associated with higher T cell activation. (A) Generation of cell lines with a modified rs11240391 variant allele by homologous recombination using a 101 pb ultramer, a single guide RNA (gRNA3), and CRISPR-Cas9 technology. Sanger sequencing chromatograms show the genomic sequence of the wild-type Jurkat clone (WT Jurkat) and of the clone in which a “T” allele has been replaced by a “G” allele (T/G Jurkat). (B) qPCR analysis of *LAX1* gene expression on wild-type clone after homologous recombination experiment (WT_HR_) and two heterozygous clones (T/G 1 and T/G 2) for the rs11240391 SNP. Values were generated in triplicate from 3 independent experiments. The plot shows the mean values ± SEM. P-values were calculated using a two-sided Student’s t-test. (C) qPCR analysis of *LAX1* gene expression in wild-type Jurkat cells (WT_HR_) and T/G clone heterozygotes (T/G 1 and T/G 2) for the SNPs rs11240391, untransfected (grey) or transfected (black) with both expression plasmids for FOS and JUN. Values were generated in triplicate from 3 independent experiments. The plot shows the mean values ± SEM. P-values were calculated using a two-sided Student’s t-test. (D) Monitoring of Jurkat cell activation by anti-CD69 staining through flow cytometry after PMA/ionomycin stimulation. The values represent the average ± SEM of two independent experiments performed in duplicate, indicating the percentage of cells positive for anti-CD69 staining (acquisition of 2000 cells per clone). The comparison was carried out between a wild-type clone after a homologous recombination experiment (WT_HR_) and two heterozygous clones (T/G 1 and T/G 2) for the rs11240391 SNP. (D) T cell activation was assessed based on CD69 staining and flow cytometric gating strategy. Plots showing the results at different time points after PMA/Ionomycin stimulation of wild-type clone after CRISPR-Cas9 editing (WT_HR_) and heterozygous clones (T/G 1 and T/G 2) for the rs11240391 SNP. Representative experiments on 2000 cells of each type in function to Forward Scatter-Horizontal (FSC-H) and anti-CD69 staining with Fluorescein Isothiocyanate (FITC). A higher percentage of CD69-positive cells was observed for T/G 1 and T/G 2 clones.

We then assessed the effect of the addition of FOS and JUN proteins in cultured wild-type cells or heterozygous clones. As expected, a significant increase in *LAX1* expression was observed in WT_HR_ (T/T) cells (p < 0.0001, 1.65-fold increase) when the 2 TFs were added together (Figure 5C). Only a slight increase in *LAX1* was observed in the presence of FOS and JUN in the T/G clones compared to the WT_HR_ (T/T) clone. This result is consistent with a lower binding of FOS and JUN to the ’G’ allele of rs11240391, as predicted by the RSAT tool.

### Heterozygous rs11240391 clones display hyperactivation of TCR signalling

Based on the biological function of the *LAX1* gene, we hypothesized that a decrease in *LAX1* expression, as demonstrated in rs11240391 heterozygous clones, may increase the activation of TCR signaling. To confirm this hypothesis, we quantified the CD69 marker on the cell surface by flow cytometry, before (0 hour) and at several points after PMA/ionomycin stimulation (at 1 hour, then every 15 minutes between 1 and 2 hours, and finally after 4 hours of stimulation) for WT_HR_ cells and the two T/G clones (Figure 5D and 5E). We observed that the number of CD69-positive cells was higher in the two heterozygous clones even before the stimulation (Figure 5D and 5E), which is consistent with the reduced *LAX1* expression of these clones. The number of CD69-positive cells was 8.3% for wild-type T/T cells (WT_HR_) compared with 28.9% and 17.1 % for the two heterozygous cells, suggesting basal hyperactivation of the cell. The number of CD69-positive cells increased with the duration of stimulation for all three clones, with higher numbers in the T/G clones than in the T/T clone (p = 0.0003 and p = 0.0008 when WT_HR_ was compared to T/C_1_ and T/C_2,_ respectively), with almost 100% of positive cells at 4 hours of stimulation for the heterozygous clones. These data demonstrate that rs11240391, a regulatory variant of *LAX1*, affects T-cell activation (Figure S1).

### Association of severe malaria with rs11240391 that could not be detected in GWAS

Since T-cell immunity plays a central role in protection against pathogens^35–37^, genetic variants that deregulate it could be associated with susceptibility to infectious diseases. Furthermore, as the *ATP2B4* internal promoter (ESpromoter) harbors genetic variants associated with severe malaria^21,37^ and regulates *LAX1* expression, we hypothesize that the rs11240391 variant in the *LAX1* promoter might also be associated with severe malaria. We conducted a case-control association study in a Senegalese population (Figure 6A, Table S1), with observed genotype frequencies consistent with Hardy–Weinberg equilibrium. We found that the GG genotype leads to an increased risk of severe malaria (OR = 2.6, p = 0.005) (Figure 6B, Figure S1) with 39% and 19% in cases and controls respectively. This result suggests that a diminished *LAX1* expression, leading to T-cell hyperactivation, may predispose individuals to disease. To assess whether *LAX1* expression might be involved in susceptibility to severe malaria, we re-analyzed previously published expression data by comparing the transcriptome of PBMCs from children with cerebral malaria (one form of severe malaria) and uncomplicated malaria^6,7^. We observed that cerebral malaria was associated with a significant reduction of *LAX1* expression (p = 0.025, fold change 1.85) (Figure 6C).

**Figure 6.**
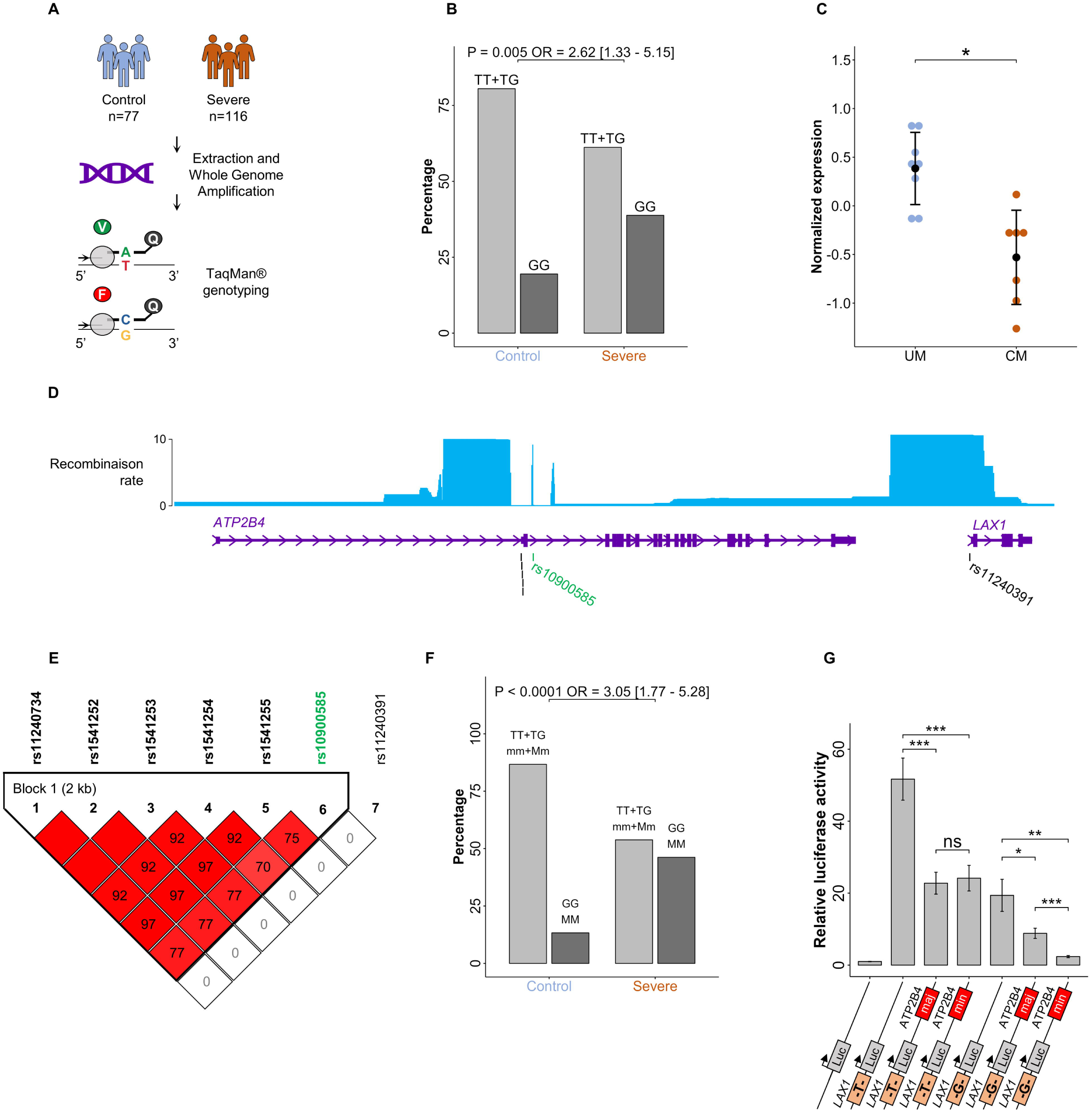
Epistatic interaction between severe malaria risk variants with decreased *LAX1* expression. (A) Design for recruitment of cohort, amplification of DNA, and genotyped of rs11240391. (B) Association of rs11240391 with severe malaria. The graph shows the percentage of the GG risk genotype (black) versus heterozygous TG and major homozygous TT (grey) in the control and severe malaria groups. P-values were calculated using logistic regression analyses and the graph displays the Odd Ratio (OR) with its 95% confidence interval. (C) Normalized *LAX1* expression obtained by microarray in children with uncomplicated malaria (UM) and cerebral malaria (CM). Lower expression of *LAX1* is observed in CM. (D) WashU epigenome browser view of recombination rate of YRI (Yoruba in Ibadan, Nigeria). Black lines correspond to the 5 SNPs previously identified (rs11240734, rs1541252, rs1541253, rs1541254, and rs1541255) and the SNP in the *LAX1* promoter (rs11240391). The green line corresponds to the tagSNP (rs10900585) previously associated with severe malaria in GWAS analyses. (E) Linkage disequilibrium (LD) between different SNPs of the *ATP2B4* and *LAX1* genes in our Senegalese cohort. LD is expressed as r^2^ multiplied by 100. SNPs with an r^2^ > 0.6 are considered in linkage disequilibrium and are colored red. (F) Epistatic interaction between the haplotype of 5 SNPs and rs11240391 computed by logistic regression. The plot shows the percentage of individuals carrying either both protective haplotype/genotype (mm+Mm/TT+TG) (m: minor haplotype, M: major haplotype) or both risk haplotype/genotype (MM/GG). Individuals carrying the GG genotype and the MM haplotype are at increased risk of severe malaria. P-values were calculated using logistic regression analyses and the graph displays the Odd Ratio (OR) with a 95% confidence interval. (G) Luciferase assays to functionally assess the epistatic effect between the haplotype including the 5 SNPs in the ESpromoter and the *LAX1* promoter SNP rs11240391. Graphs showing the relative luciferase activity under the control of *LAX1* promoter with “T” or “G” allele at rs11210391 alone or in combination with the ESpromoter containing either the major haplotype (maj: TCCGA) or minor haplotype (min: CTTGG) for the 5 SNPs (rs11240734, rs1541252, rs1541253, rs1541254, and rs1541255). Values were generated in 84triplicate from 3 independent experiments. The plot shows the mean values ± SEM. P-values were calculated using a two-sided Student’s t-test.

**Figure 7.**
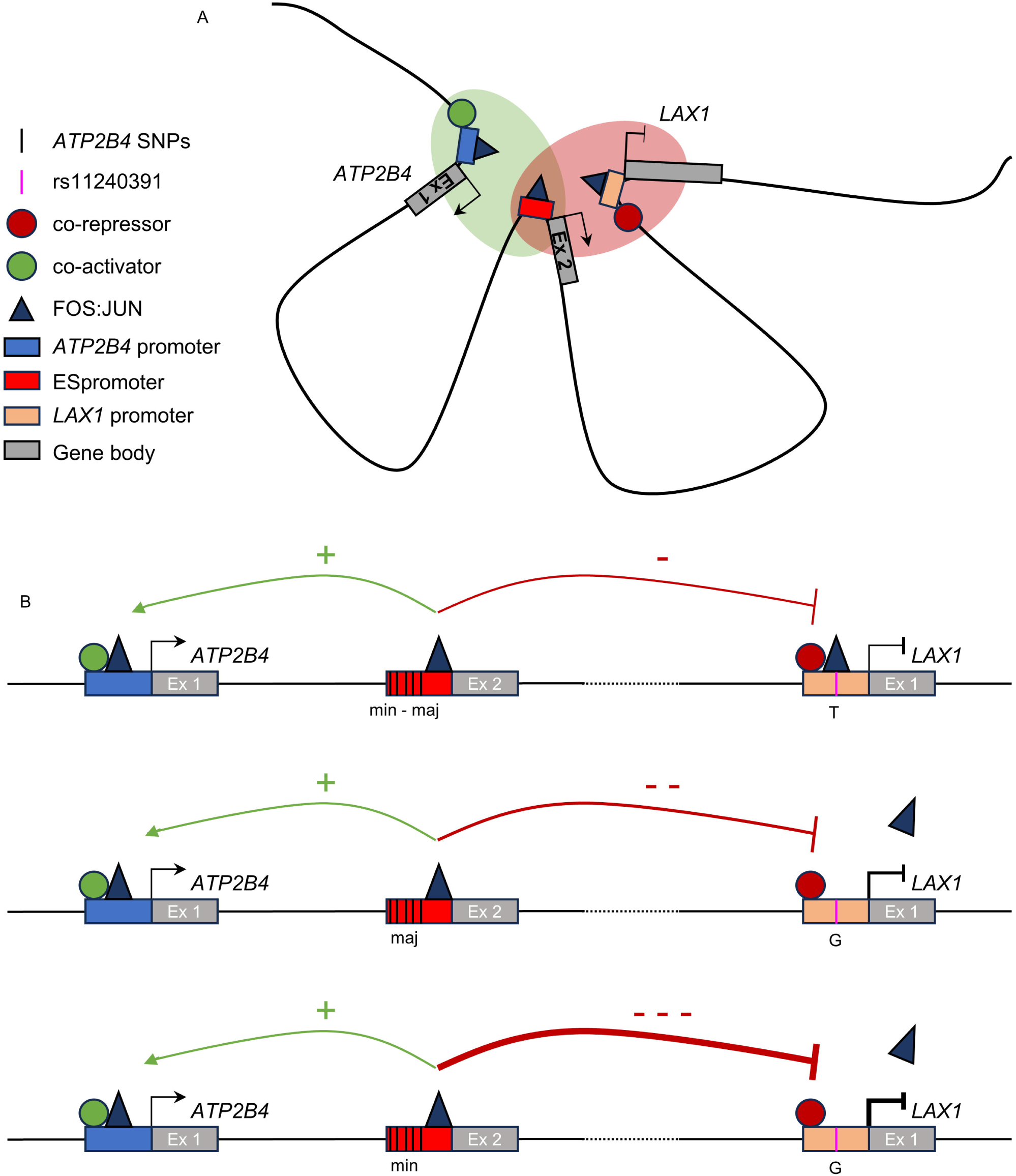
Model of dual enhancer-silencer function of ESpromoter, epistatic interaction, and gene regulation. (A) Chromatin interactions place promoters in close physical proximity, facilitating the recruitment of transcription factors needed to activate or repress transcription of their associated genes. The presence of an Enhancer-Silencer promoter (ESpromoter) within regulated genes cluster could facilitate the assembly or maintenance of the transcription factor and co-factors by tightening the promoter-promoter interactions or by providing specific transcriptional regulators required for neighboring gene regulation. (B) Expression of the long transcripts *ATP2B4* and *LAX1* is regulated by a dual enhancer-silencer regulatory element (ESpromoter) in the same cell line through genetic variants associated with severe malaria. ESpromoter functions as an enhancer for the long *ATP2B4* transcripts independently of the haplotype of the five SNPs it contains (min: minor haplotype or Maj: major haplotype) while it functions as a silencer for *LAX1* gene with an allele-specific intensity. The five SNPs within the ESpromoter act synergistically with rs11240391 to inhibit *LAX1* gene expression. This epistatic interaction results in a stronger silencing effect in the presence of the G allele of rs11240391, for which no FOS:JUN binding is possible. Unidentified co-activators and co-repressors are also thought to modulate the activating or repressing effect on *ATP2B4* and *LAX1* respectively.

It is important to note that the association signal for rs11240391, newly identified here, could not be detected in previous genome-wide association studies, as this SNP is not included among the genotyped variants and is in very low LD with all other SNPs (r^2^ < 0.4, except for rs6680886 having an r^2^ of 0.56) in African populations. The LD between rs11240391 and the TagSNP rs10900585 associated with severe malaria by GWAS was estimated to be r^2^ = 0.0015 in African populations using the LD pair module in LDlink web-based application^38^. Consistently, figure 6D shows that the SNP rs11240391 is located in a region with a high recombination rate (data from Yoruba in Ibadan, Nigeria), which favors the absence of polymorphisms in LD with it. We then confirmed the absence of LD in our Senegalese population between rs11240391 and the TagSNP rs10900585 genotyped in the GWAS, as well as with the 5 SNPs within the *ATP2B4* ESpromoter (Figure 6E).

### Epistatic interaction between rs11240391 and SNPs at ESpromoter

Considering the established involvement of the ESpromoter in *LAX1* gene regulation and the physical interaction between it and the *LAX1* promoter, we decided to explore potential epistatic interactions between rs11240391 and the five SNPs of the ESpromoter previously associated with severe malaria^17^. Individuals carrying both the GG risk genotype at rs11240391 and the *ATP2B4* risk haplotype (Major haplotype) had a higher risk of developing severe malaria with 21% and 8% in severe malaria and controls, respectively (OR = 3.05, p < 0.0001) (Figure 6F, Figure S1, Table S2), suggesting an epistatic interaction between them. We then assessed the functional relevance of this epistatic interaction by luciferase reporter assays. We transfected Jurkat cells with constructs containing the 810-bp promoter region of

*LAX1* with the T or G allele for rs11240391 upstream of luciferase, and the ESpromoter with the major or minor haplotype downstream of luciferase. We validated the haplotype-independent silencer impact of the ESpromoter on the *LAX1* promoter in the presence of the major T allele of rs11240391 (p < 0.001, with a 2.27- and 2.13-fold decrease in the presence of the major haplotype or the minor haplotype respectively).

However, in the presence of the minor G allele of rs11240391 at the *LAX1* promoter, the silencer effect of the ESpromoter was haplotype-dependent showing a more pronounced decrease in luciferase expression in the presence of the minor haplotype (p < 0.01, 2.19-fold decrease in presence of minor G allele with major haplotype versus 8.18 with minor haplotype) (Figure 6G).

We have demonstrated an epistatic interaction since the regulatory impact of the genetic variants at the ESpromoter depends on the allele of rs11240391 at the *LAX1* promoter. This underlies a functional relationship between them, in favor of a synergistic effect.

## DISCUSSION

To elucidate the regulatory dynamics of the *ATP2B4* locus and its potential impact on the risk of severe malaria, we used an integrative approach combining genetics, epigenomics, and CRISPR-cas9 genome editing. We aimed to decipher the molecular mechanisms by which regulatory elements and causal variants influence gene function. We discovered a hitherto unknown type of cis-regulatory element, which we called “ESpromoter”. To our knowledge, this is the first description of a promoter element with dual enhancer and silencer functions that simultaneously activates and represses gene expression in a single human cell type. Our strategy also enabled us to identify a new functional SNP (rs11240391) associated with severe malaria, raising a fundamental point about the weakness of genome-wide association analyses for identifying causal variants in complex human diseases.

Transcriptional enhancers play a crucial role in gene regulation by activating gene expression in a tissue-specific manner during development and in adult cells in response to cellular signals or environmental stimuli. However, it is also important that gene expression is not inappropriately activated or regulated. Transcriptional silencers are active negative regulatory elements that repress promoter transcription^39^. They play a crucial role in helping to specify precise patterns of gene expression by preventing ectopic expression. Interestingly, some regulatory elements have previously been found with both functions, which can act either as an enhancer or a silencer, depending on the type of tissue or cellular context, in Drosophila^40–42^, in mice^43^ and human^44^. Recently, it has been suggested that most silencers are also enhancers in different cell types in Drosophila^41^ and also that promoters may have silencer-like function^45,46^. However, to our knowledge, no element with the simultaneous dual enhancer-silencer function in the same cell type has been discovered in humans.

Our results therefore challenge the current view of enhancers and silencers as two distinct types of regulatory elements. Instead, we suggest that dual-function CREs in the same cell type may be an important phenomenon in transcriptional regulation that has not yet been explored reflecting the complexity, dynamics, and diversity of regulatory interactions at active elements. Deletion of the ESpromoter in Jurkat T cells resulted in both an increase in *LAX1* and a decrease in *ATP2B4* expression, demonstrating the importance of this element for the proper expression level of its target genes. Currently, the exact mechanism underlying these dual elements in the same cell type remains unknown. However, there is supporting evidence for a model of enhancer-silencer activity in which the ESpromoter would make direct 3D contact with the promoters of the regulated genes^27^, likely facilitated by TFs and cofactors acting as intermediaries. Cis-regulatory elements are DNA fragments that serve as active hubs to recruit TFs through short DNA motifs to regulate transcription. TFs binding and its impact on gene expression are influenced by many different mechanisms, ensuring robust and complex spatio-temporal regulation^47^. In particular, the positioning of TFs appropriately facilitates protein-protein interaction and thus promotes cooperative binding as well as the recruitment of cofactors and the transcriptional machinery^47^. Here, we demonstrated that *LAX1* expression required both FOS and JUN transcription factors, suggesting that this co-recruitment may represent cooperative binding to the *LAX1* promoter. This cooperative binding is efficient only in the presence of the major allele T at rs11240391. The Activator Protein 1 (AP-1) a dimeric complex consisting of FOS and JUN, may regulate gene expression in response to stimuli such as cytokines and infection and has been described as a pioneer TF allowing chromatin remodelling. Interestingly, AP-1 is also able to bind to the ESpromoter region (Figure S3) independently of the haplotype, but also to the *ATP2B4* promoter in which no common variants were identified. However, we do not know whether AP-1 binding in the 3 regions could play a direct role in the dual enhancer-silencer function of the ESpromoter. Indeed, it has been described that specific TFs, such as GATA3, can function either as activators for particular genes or as transcriptional repressors for others, acting directly on the transcriptional machinery and/or affecting chromatin remodeling at crucial loci, particularly those implicated in Th1/Th2 differentiation^48^.

We can also hypothesize that TFs at the ESpromoter act cooperatively with specific TFs on either the *ATP2B4* or *LAX1* promoter, leading to activation or repression of these genes respectively, as illustrated in Figure 6. Some TFs were reported to have repressive activity, such as REST, YY1^49^ while others can act as activators^50^. In some cases, cis-regulatory elements contain genetic variants, adding further complexity to their genomic function. Up to 90% of GWAS loci contain non-coding variants and the precise identification of disease-causing regulatory variants within GWAS loci remains a major challenge, particularly in terms of prioritization and validation of the putative functional effects of these variants. Here, we have successfully identified a novel regulatory SNP associated with severe malaria, rs11240391 in the *LAX1* promoter. This SNP undetectable by conventional GWAS methods due to its non-inclusion as a TagSNP and its lack of LD with other SNPs in African populations. Although GWAS have made it possible to localize loci associated with complex diseases, relying solely on these studies presents limitations in identifying causal functional SNPs and requires an alternative strategy. Our multi-approaches thus pave the way for a new strategy for discovering causal variants. Furthermore, such causal regulatory variants, which often reach a high frequency in populations, individually have a low impact on gene expression and are likely to affect disease risk through modest phenotypic effects. We, therefore, propose that functional assessment of common non-coding variants is often more relevant when several variants are tested simultaneously in order to assess their epistatic interaction and functional consequences^17,21^. Here, we found that the enhancer or silencer function is independent of the *ATP2B4* haplotype in the presence of the major T allele at rs11240391 in Jurkat T cells, whereas the repressive function of the ESpromoter is stronger in combination with the minor G allele for rs11240391, suggesting an epistatic interaction between all these SNPs. This is consistent with the genetic interaction that we observed between these variants, resulting in a significant increase in the risk of severe malaria in the Senegalese population. In addition, common variants located in dual enhancer-silencer elements can synergistically or antagonistically modify the regulation of several nearby genes, which may be involved in the same biological pathway, potentially leading to stronger molecular perturbations and important phenotypic consequences. Our work provides a general approach to improve the identification of novel regulatory variants associated with complex diseases. We and others^51^ have demonstrated that integrating information from GWAS with epigenomic features is effective in discovering novel regulatory elements and polymorphisms missed by GWAS for various reasons (low linkage disequilibrium, sub-threshold signal, cell-specific effect).

Interestingly, our results confirmed that this ESpromoter regulates the *ATP2B4* and *LAX1* genes, which respectively code for plasma membrane Ca2+ ATPase (PMCA) and the adaptor protein LAX, two proteins playing an essential role in T-cell activation^30,31^. PMCA, encoded by 4 different genes (*ATP2B1-4*), is the main extrusion pump expressed in various cell types regulating intracellular Ca2+ homeostasis. Analysis of the expressed isoform of PMCA revealed that PMCA4 (encoded by *ATP2B4*) is the most predominant and functional in CD4+ T cells^30^. Specifically, PMCA4b (transcript ENST00000357681), one of the two long transcripts regulated by the ESpromoter, is the major transcript in CD4+ T cells and Jurkat cells^52^, which is also more highly expressed in memory cells than in naive cells^30^. As the downregulation of PMCA4 reduced the effector cell fraction and led to the accumulation of naïve cells^30^, the dual cis-regulatory element identified here may play a pivotal role during CD4+ T cell activation by regulating the stoichiometry of CD4+ T cell subpopulation and, consequently the outcome of the immune response. As previously suggested by others, transcriptional regulation of PMCA4 is likely to be biphasic^30^, but we propose a different scenario in which the increase in PMCA4 expression is mediated by the enhancer function of the ESpromoter. Firstly, when CD4+ T cells are activated, YY1 is upregulated and represses PMCA expression so that CD4+ T cells can adapt to higher intracellular levels of cytosolic Ca2+ to support gene expression and proliferation. Secondly, PMCA4 expression increases to protect against intracellular Ca2+ overload and promote the survival of effector cells and the formation of long-lasting memory cells capable of efficiently mediating a more rapid immune response upon subsequent encounter with an antigen. In addition, CD147 has been shown to interact via its immunomodulatory domains with PMCA4 to bypass proximal TCR signaling and inhibit IL-2 expression^53^.

CD4+ T cell differentiation or regulation is a complex process in which a plethora of genes are tightly orchestrated, and *LAX1* appears to be one of these genes in addition to *ATP2B4*, confirming that the ESpromoter is a key regulatory element in this process. The LAX adaptor protein, an LAT-like molecule identified in T cells^54^ functions differently from LAT to negatively regulate the TCR signaling pathway^31^. Consistently, we demonstrated that deletion of the ESpromoter was associated with overexpression of *LAX1* leading to a decrease in the number of CD69-positive cells upon stimulation. Furthermore, Jurkat clones heterozygous at rs11240391, in which *LAX1* was downregulated, were hyperresponsive to TCR-mediated stimulation in a similar manner to LAX-deficient mice^55^. We, therefore, provide further evidence for the role of LAX as a negative regulator of lymphocyte signaling, acting probably by inhibiting TCR-mediated p38 MAPK and NFAT/AP-1 activation, as previously proposed^54^. Interestingly, calcium flux is also enhanced in LAX-deficient mice. Overall, our data suggest a model, in which PMCA4 and LAX, whose *ATP2B4* and *LAX1* genes are regulated by a dual enhancer-silencer element, play a central role in the formation of Ca2+ signals in different states of CD4+ T cell activation. This suggests that several positive and negative regulatory mechanisms work together to fine-tune TCR signaling, aiming for a balanced immune response.

Moreover, ESpromoters may have pleiotropic impacts on various physiological and pathological traits, by differentially regulating multiple proximal and distal genes at a given locus^18^. These genes may not necessarily be part of the same biological pathway but could be involved in pathogenic processes shared across different diseases. Consequently, dysfunction of cis-regulatory elements due to genetic, structural or epigenetic alterations may cause a wide range of human diseases^56^. It would therefore not be surprising if genetic variations within the ESpromoter (rs11240734, rs1541252, rs1541253, rs1541254, and rs1541255) and the *LAX1* promoter (rs11240391) could have pleiotropic effects, since they regulate the function of T cells, an important mediator of the adaptive immune system, capable carrying out coordinated and specific responses to combat infection and prevent inflammation. The pleiotropic role of non-coding variants has recently been pointed up by the identification of a genetic variant associated with five vascular diseases^57^. Finally, we have highlighted another layer of complexity by demonstrating an epistatic interaction between these genetic variants, which increases the risk of severe malaria, leading to greater dysregulation of *LAX1* with hyperactivation of T cells. Understanding epistasis is key to unraveling the complexity inherent in genetic combinations and their effect in complex diseases.

Although we do not yet have evidence of a 3D interaction between the ESpromoter and the promoter of the target genes in CD8+ T cell, open chromatin can clearly be observed for the *ATP2B4* and *LAX1* promoters as well as for the ESpromoter, suggesting a similar regulatory mechanism in this cell type. We can therefore speculate that hyperactivation in CD8+ T cells following deregulation of *LAX1* and deregulation of intracellular Ca2+ signaling, a major determinant of CD8+ T cell reactivity, may also play an important role in determining severe malaria and more particularly cerebral malaria. In addition, hyperactivation of the immune system lies at the center of many autoimmune and allergic diseases and it is prevented by upregulation of immunosuppressive proteins following lymphocyte activation^53^. Hence, identifying the regulatory elements involved in the regulation of the immune response is crucial.

To our knowledge, we were able to demonstrate for the first time the contribution of a novel regulatory element, the ESpromoter, a promoter acting as both enhancer and silencer in a particular cell type. More specifically, this dual element simultaneously up-regulates *ATP2B4* gene and down-regulates *LAX1* gene. Their expression is modulated by epistatic interaction of genetic variants within the ESpromoter and the *LAX1* promoter, leading to their deregulation and subsequently potential impairment of T cell functionality. We suggested that the widely observed notion that silencers constitute a distinct class of regulatory elements from enhancers is an oversimplification and that dual function may be a common phenomenon in transcriptional regulation. Additionally, some variants of regulatory elements may impact the risk of several diseases exhibiting a pleiotropic effect. Finally, a combined analysis of genetic and epigenetic characteristics is necessary to allow the identification of the causal variants not found by GWAS alone.

## SUPPLEMENTAL INFORMATION

Supplemental Information includes 3 figures and 3 tables that can be found with the article online.

## ACKNOWLEDGMENTS

This work was supported by the African Higher Education Centers of Excellence project (CEA-SAMEF) at UCAD (to B.M.); the Pasteur Institutes in Dakar and in Paris (to A.T.); the French Embassy in Dakar; recurrent funding from Institute National de la Santé et de la Recherche Médicale (INSERM) and Aix-Marseille University; the French Agency for Research (Agence Nationale de la Recherche ANR) MADBIO ANR-22-CE15-0025-01 (to S.M.). M.A. and S.N. were supported by a PhD fellowship from the French Ministry of Research and the Higher Education Commission (HEC) from Pakistan, respectively. This project was carried out in the framework of the INSERM GOLD Cross-Cutting program (P.R., S.M.). We thank the members of TAGC, Lydie Pradel and Nathalie Arquier for helpful discussions and Pauline Andrieux for her advice on CRISPR technology. Selected artwork shown in the graphical abstract were used from or adapted from pictures provided by Servier Medical Art (Servier; https://smart.servier.com/), licensed under a Creative Commons Attribution 4.0 Unported License. We thank all patients who participated in the study, the Dakar main hospital and Tambacounda regional hospital for the recruitment of biological samples.

## AUTHOR CONTRIBUTIONS

Conceptualization, S.M.; Methodology, M.A., S.N., A.E., H.T.N.H., M.T., and I.M.; Investigation, S.S., P.R., and S.M.; Senegalese cohort, A.T., B.M., A.D., and P.R.; Writing - Original Draft, M.A., and S.M.; Writing - review and Editing, M.A., S.S., P.R., and S.M., Funding Acquisition, A.T., A.D., P.R., and S.M.; Supervision, S.M.

## DECLARATION OF INTERESTS

All authors declare no competing interests.

## STAR METHODS

**KEY RESSOURCES TABLE**

## RESOURCE AVAILABILITY

### Lead contact

Further information and requests for resources and reagents should be directed to and will be fulfilled by the Lead contact, Sandrine Marquet.

### Experimental Model and Subject Details Cell lines

Jurkat cells were cultured according to standard cell culture protocols. RPMI 1640 medium with GlutaMax (61870-044, ThermoFisher, Waltham, MA, USA) supplemented with 10% fetal bovine serum (FBS, A5256701, ThermoFisher, Waltham, MA, USA) was used as the culture medium. Cells were maintained in a 37 °C incubator with 95% humidity and an atmosphere containing 5% CO2. Jurkat cells were subcultured to a density of 1 million cells per milliliter. The culture medium was refreshed every three days to ensure a constant supply of nutrients. To activate Jurkat cells, combined stimulation with phorbol 12-myristate 13-acetate (PMA; Sigma-Aldrich, P1585) and ionomycin (Iono; Sigma-Aldrich, I3909) was performed. Cells were harvested during the exponential growth phase, washed, and resuspended in a fresh medium. Subsequently, PMA, a protein kinase C activator, and ionomycin, a calcium ionophore were added at 20 ng/mL and 2.5 µM final final, respectively. The cells were incubated for a specific time in each experiment.

### Cohort

Patients diagnosed with malaria were recruited from two Senegalese locations, Dakar Primary Hospital and Tambacounda Regional Hospital, including a cohort of 90 cases of cerebral malaria and 27 cases of severe non-cerebral malaria, as reported previously^58^. Control samples (n = 79) were recruited from healthy volunteers in Dakar. Written informed consent was obtained from each patient and accompanying family members. The research protocol was approved by the institutional research ethics committee of Cheikh Anta Diop University.

On the day of admission, venous blood samples were collected alongside biological data, to determine parasite density, haematology, and other relevant characteristics. The presence of *Plasmodium falciparum* was checked by examination of thin and thick blood smears by at least two biologists trained before initiation of antimalarial treatment. A blood smear was considered positive if asexual parasites were identified, and quantification of parasitized erythrocytes was performed by counting the number per microliter (µL) of blood.

### Method Details Data acquisition

The data of physical chromatin interactions were obtained from Genehancer data prediction^59^ and from GEO: GSM6509371^27^. The circos plot was built using ClicO FS^60^. All genomic regions were obtained using WashU Epigenome Browser^61^. ATAC-seq data from a PBMC sub-cell were obtained from Calderon et al.^28^. Naive CD4+ epigenomic markers were obtained from Roadmap Data from GEO available on WashU Epigenome Browser^29^. ReMap data^34^ were obtained from the download page (https://remap.univ-amu.fr/download_page) using non-redundant peaks. The predicted regulatory elements come from the ENCODE project^62^ available on UCSC tracks. The recombination rate data is also available on the UCSC website and comes from the HapMap project^63^. The eQTL data and pip value were obtained from ELIXIR Estonia eQTL Browser^32^.

### Motif analysis

The impact of the selected SNP on transcription factor binding sites (TFBS) was assessed with the Rsat tool^64^, in its default settings and using the Jaspar TFBS databases^65^ (Jaspar core non-redundant vertebrates 2020 Collection). Then, for the predicted transcription factor binding sites, the presence of transcription factor in the SNP region was confirmed using ReMap data^34^.

### Genome editing using CRISPR-Cas9

#### CRISPR/Cas9 guide selection, single-stranded oligodeoxynucleotide donor design, and RNP complex preparation

We identified guide RNAs using computational algorithms prioritizing on-target efficacy and reduction of off-target effects using the CRISPR design tool provided by IDT (Integrated DNA Technologies), the custom Alt-R^TM^ CRISPR-Cas9 guide RNA and the Alt-R^TM^ CRISPR HDR design tools. A 101-bp HDR donor oligonucleotide was also designed for CRISPR-mediated Homology Directed Repair (HDR). These chemically synthesized oligoribonucleotides were manufactured by IDT: crRNAs (35 mer with a part specific to the target DNA sequence, Alt-R^®^ CRISPR-Cas9 crRNA, IDT), universal tracrRNAs (67 mer, Alt-R CRISPR-Cas9 tracrRNA, 20 nmol, 1072533, IDT) and the 100-bp single-stranded oligodeoxynucleotide donor template (ssODN). These sequences were resuspended in IDTE buffer (pH 7.5, 11-05-01-15, IDT) to achieve a final concentration of 100 μM each. The active gRNA complex was formed by mixing 5 μL of universal tracrRNA (100 μM) with 5 μL of crRNA (100 μM) and incubated for 5 minutes at 95°C and then returned to room temperature. The Cas9 Ribonucleoprotein (RNP) complex was assembled *in vitro* by incubating 3.4 µL of Cas9 protein (62 µM) (Alt-R^TM^-S.p Hifi Cas9 Nuclease V3, 100 µg, 1081060, IDT) with 4.8 µL active gRNA complex (crRNA-tracrRNA) and 1.8 µL of PBS (1X).

### Deletion of ESpromoter region

The ESpromoter deletion was performed in Jurkat cells as previously described^17^. To generate the 506 bp genomic deletion comprising the 5 SNPs, 5 x 10^5^ Jurkat cells were electroporated by the Neon transfection system (MPK5000, Invitrogen) with 2 µL of gRNA1 RNP complex and 2 µL gRNA2 RNP complex and 2 μL of HDR enhancer. Following transfection, the pool of transfected cells was clonally plated into 96-well plates at a limiting dilution of less than 0.5 to avoid mixed clones. Following 14 days of cell growth, the individual clone was isolated, genomic DNA was extracted and amplified by PCR, then screened for the desired deletion. DNA amplification was performed directly from the clones, previously treated with 20 µL of dilution buffer and 0.5 µL of DNA Release additive, which improved DNA release from the cells. Polymerase chain reaction (PCR) was performed using 25 µL of 2X Phire Tissue Direct PCR Master Mix containing Phire Hot Start II DNA Polymerase (Thermo Fisher Scientific, Waltham, MA, USA), 2 µL of previously prepared DNA and 1 µL of forward and reverse primer (10 µM), annealing at 60 °C. Clones were screened for deletion by PCR using primers F1 and R1, followed by agarose gel electrophoresis of PCR amplicons. A short fragment of 335 bp in the presence of deletion and a long fragment of 841 bp in the absence of deletion were expected. PCR products were purified using the PCR DNA Purification kit (QIAGEN), according to the manufacturer’s instructions, and the deletion was confirmed by Sanger sequencing of PCR amplicons. One wild-type clone (WT_c_) and three 506-bp biallelic deletion clones (Δ1, Δ2, and Δ3) were selected for further study.

### Single nucleotide editing in Jurkat cells

The Jurkat cell line was genotyped for rs11240391 by PCR amplification of a 527 bp fragment using primers F2 and R2, followed by Sanger sequencing. The sequence indicated that the Jurkat cell line is homozygous T/T for rs11240391. To replace, by homologous recombination, the T allele with a G allele at the rs11240391 variant located in the promoter region of the *LAX1* gene, we transfected the cells with 3.6 μL of gRNA3 RNP complex, 0.4 μL HDR enhancer (3mM) and 3.6 μL at 100 μM of ssODN in which a G allele is present in place of the T allele. The active gRNA complex and Cas9 RNP complex were made as described above and transfected by the Neon transfection system into Jurkat cells. After 48 hours, the pool of transfected cells was clonally plated into 96-well plates at a limiting dilution of less than 0.5 to avoid mixed clones. Cell growth, DNA extraction, and PCR amplification using primers F2 and R2 were performed as described above. The sequence of the rs11240391 variant was determined by Sanger sequencing in different clones. One wild-type clone (WT_HR_) and two heterozygous clones (T/G_1_ and T/G_2_) were selected for further study.

#### sgRNA sequences

gRNA1: TCCTCTACATTGGAGTTTAC AGG gRNA2: TAGACTTCGGACGGCTACTC GGG gRNA3 : CCAATGTGCTAATGAAGCAC AGG

#### ssODN sequence

5’-TATGTTTTCTTCTAGCAGATTAAGAGCTGAGCAGAGTTTCCTGTGCCCTGGGCTTCATTAGCACATTGGTGGTGTCGTTTCCGGTGACTGACTCTCTGTTT-3’

#### PCR primers

F1: primer forward, GGCCACCCTTCAGATCACTT R1: primer reverse, GCCTCCCTGTCTCAACTTCT F2: primer forward, TGAATCAGAAGAGGGTCCCG R2: primer reverse, CGATCTCACCGGACATGGT

### Luciferase reporter assay Promoter Activity

The 780-bp *ATP2B4* promoter fragment (GRCh38, chr1: 203626081-203626860) was inserted into the MlulI-XhoI sites of the pGL3-basic vector (Promega, Madison, WI, USA, # E1751), which contained the firefly luciferase coding sequence (GeneCust, Boynes, France). The 810-bp *LAX1* promoter region (GRCh38, chr1: 203764726-203765536) containing rs1124039 was also inserted into the MlulI-XhoI sites of the pGL3-basic vector (Promega, Madison, WI, USA, # E1751). These constructs were supplied by Genecust custom services (Luxembourg). Site-directed mutagenesis was then performed to modify the “T” allele to “G” in rs1124039 using the Q5 Site-Directed Mutagenesis Kit (New England Biolabs) and the Forward 5’ – CTGTGCCCTGgGCTTCATTAG -3’ and Reverse 5’ – GAAACTCTGCTCAGCTCTTAATC - 3’ mutagenesis primers designed by NEBaseChanger tool.

Jurkat cells were transfected using the Neon^TM^ Transfection system (Invitrogen) according to the manufacturer’s instructions. One day prior to transfection, cells were diluted to a concentration of 0.6 million. In each experiment, 1 million cells were co-transfected with 1 µg of vectors: (1) negative control vector (empty pGL3-basic vector (cat# E1751)) or the construct to be tested (pGL3-ATP2B4 promoter, pGL3-LAX1 rs11240391-T allele, pGL3-LAX1 rs11240391-G allele) and (2) 200 ng of pRL-SV40 (plasmid encoding renilla luciferase from Promega (cat# E2231)), as an internal control for transfection efficiency. After transfection, Jurkat cells were rested at 37°C in 5% CO2 for 24 h. Under stimulation conditions, Jurkat cells were treated with PMA/ionomycin and incubated for 6 hours or 24 hours. Next, cells were subjected to firefly and renilla luciferase activity on 20 µl of cell lysate with 100μL LARII (1X) followed by 100μL of Stop and go (1X) according to the standard instructions provided in the Dual-Luciferase kit (Promega, Madison, WI, USA) using a TriStar LB 941 Multimode Microplate Reader (Berthold Technologies, Thermo Fisher Scientific, Waltham, MA, USA). The firefly luciferase activity of each sample was normalized to Renilla luciferase and expressed as fold change relative to the empty vector control. Three experiments were performed in triplicate.

### Enhancer-silencer Activity

From the construct containing the 780-bp fragment of *ATP2B4* promoter (GRCh38, chr1: 203626081-203626860) at the MluI-XhoI sites, the 601-bp fragment of *ATP2B4* ESpromoter (GRCh38, chr1: 203682499-203683100) was cloned into the SalI-BamHI sites either with the minor or major alleles of the five SNPs (rs11240734, rs1541252, rs1541253, rs1541254, and rs1541255). The 601-bp fragment of *ATP2B4* ESpromoter containing the respective minor or major allele of the 5 SNPs was cloned into the SalI-BamHI sites in the construct having the 810-bp of LAX1 promoter region (GRCh38, chr1: 203764726-203765536) at the MluI-XhoI used for the promoter activity. Luciferase assays were performed in Jurkat cells as described above.

### Effect of transcription factors

For the luciferase assay, the cells were transfected with a total of 2 µg of plasmid, including (1) 0.5 μg of pRL-SV40 and (2) 0.5 μg of either Firefly luciferase gene pGL3-LAX1 rs11240391-T allele or pGL3-LAX1 rs11240391-G allele with or without (3) 0.5 μg of expression plasmids coding for proteins of interest (FOS, RC202597, Origene) and or (JUN, RC209804, Origene) completed with (4) 0 to 1 μg of empty pcDNA3.1 plasmid to equalize DNA quantities as previously described^66^ in each condition as detailed below: 1: 0.5 μg pGL3-Renilla + 0.5 μg pGL3-LAX1-T + 1 μg pcDNA3.1 2: 0.5 μg pGL3-Renilla + 0.5 μg pGL3-LAX1-G + 1 μg pcDNA3.1 3: 0.5 μg pGL3-Renilla + 0.5 μg pGL3-LAX1-T + 0.5 μg plasmide JUN + 0.5 μg pcDNA3.1 4: 0.5 μg pGL3-Renilla + 0.5 μg pGL3-LAX1-G + 0.5 μg plasmide JUN + 0.5 μg pcDNA3.1 5: 0.5 μg pGL3-Renilla + 0.5 μg pGL3-LAX1-T + 0.5 μg plasmide FOS + 0.5 μg pcDNA3.1 6: 0.5 μg pGL3-Renilla + 0.5 μg pGL3-LAX1-G + 0.5 μg plasmide FOS + 0.5 μg pcDNA3.1 7: 0.5 μg pGL3-Renilla + 0.5 μg pGL3-LAX1-T + 0.5 μg plasmide JUN + 0.5 μg plasmide FOS 8: 0.5 μg pGL3-Renilla + 0.5 μg pGL3-LAX1-G + 0.5 μg plasmide JUN + 0.5 μg plasmide FOS Luciferase assays were performed in Jurkat cells as described above.

### Transfection of WT and T/G edited clones with transcription factors

Jurkat WT clone WT_HR_ (wild-type clone isolated after CRISPR/Cas 9) and heterozygous clones T/G_1_ and T/G_2_ were counted before being diluted to 0.3 million cells per milliliter in RPMI+glutamax culture medium (10% decomplemented SVF) 24 h before transfections. Cells were washed in 10 mL of 1X DPBS. For each clone, 1 million cells resuspended in 100 µL of T buffer (NEON Invitrogen) were co-transfected with 500 ng of FOS and 500 ng of JUN expression plasmids (RC202597 and RC209804 respectively, Origene). In the condition without plasmids, the clones underwent the electric shock of transfection (1200 volts 40 ms 1 pulse) and will serve as controls. For each sample, each condition was performed in two replicates. Cell pellets were recovered after 48h of cell culture.

### RT-qPCR

RNA extractions from Jurkat wild-type cells and edited clones (deletion or allele modification) were conducted using the RNeasy Plus mini kit (Qiagen, Hilden, Germany). 1 µg of RNA per sample was transcribed into cDNA using the Superscript VILO Master Mix (Invitrogen, Thermo Fisher Scientific, Waltham, MA, USA). Real-time quantitative PCR (RT-qPCR) was performed using the SYBR Select Master Mix (ThermoFisher Scientific, Applied Biosystems Waltham, MA, USA) on the QuantStudio 6 Flex instrument on 10-fold diluted cDNA. Primers F3/R3 for *ATP2B4* and F4/R4 for *LAX1* gene expression quantification were designed using Primer3 software^67^. Gene expression was normalized using HPRT1 housekeeping gene, and the relative expression was computed using the ΔCT method. The data provided is an average of triplicates from three independent experiments per sample.

#### PCR primers

F3: primer forward, CAAGAGTCTGGCCCGAGTTA R3: primer reverse, TCTGCTGTTGAGATCGTCCA F4: primer forward, AGAAATTCTGAGAGCCCGGAG R4: primer reverse, GATACCCACCGCGTACTCTG

### Flow cytometry

The day before PMA/Ionomycin stimulation, cells were diluted to a concentration of 0.6 million. The cells were then stimulated as described above. After stimulation, samples were collected at 6 time points: 0h, every 15 minutes from 1h to 2h, and a final point at 4 h. Samples were stored at 4°C until collection of the last time point at 4 h. For each time point, samples were divided into two groups: unstained cells (negative) and stained cells (positive) with an anti-CD69 antibody (310904, BioLegend). The cells were then centrifuged at 1500 rpm for 5 minutes at 4°C. Positive samples were resuspended with 100 µL of anti-CD69/DPBS antibody. After 30 minutes of incubation in the dark at 4°C, samples were washed with 3ml DPBS and centrifuged at 1500 rpm. Cells were then resuspended in 250 µL of DPBS. Negative samples were resuspended in 250 µL of DPBS. Samples were acquired on the LSRFortessa-X20 cytometer (BD Bio-sciences), and data were analysed using FlowJo software (version 10, LLC).

### TaqMan genotyping

Genomic DNA from the Senegalese population was extracted and amplified as described previously^58^. The rs11240391 single nucleotide polymorphism (SNP) was genotyped using the TaqMan allelic discrimination technique (4351379, C_319219_10, Thermofisher, Waltham, MA, USA) on the QuantStudio 6 Flex instrument. A total of 117 severe malaria cases and 79 controls were genotyped. The master mix consisted of 1 µl of genomic DNA at a concentration of 12.5 ng/µl, 2.1 µl of 2x master mix (TaqMan Genotyping Master Mix, Applied Biosystems, Waltham, MA, USA), and 0.06 µl for the Taqman 40x assays in a final volume of 5 µl. The thermal cycling program included an initial step of 10 minutes at 95°C, followed by 40 cycles.

### Statistical analyses

The Haploview tool^68^ was used to assess the deviation of genotype frequency from Hardy Weinberg equilibrium in the control group. Genetic association analyses were conducted using IBM SPSS Statistics 27 (IBM, NY, USA). Logistic regression analyses were performed to adjust for the influence of age and to determine genetic interactions between *ATP2B4* and *LAX1* gene variants. Statistical analyses of luciferase reporter assays and RT-qPCR results were performed by Student’s t-tests (ns, * p < 0.05, ** p < 0.01, *** p < 0.001, **** p < 0.0001). All tests were two-tailed. Graphs were generated using ggplot2^69^ and ggpubr. Paired t test was used to compare the percentage of CD69-positive cells of wild-type cells with that of cells mutated with the Crispr-Cas9 method, after assessing the normality of the data using Shapiro-Wilk method. These tests were one-sided, with an alternative hypothesis based on known *LAX1* expression in the cells and the known effect of *LAX1* on CD69 expression.

## SUPPLEMENTAL INFORMATION TITLES AND LEGENDS

**Figure S1.**
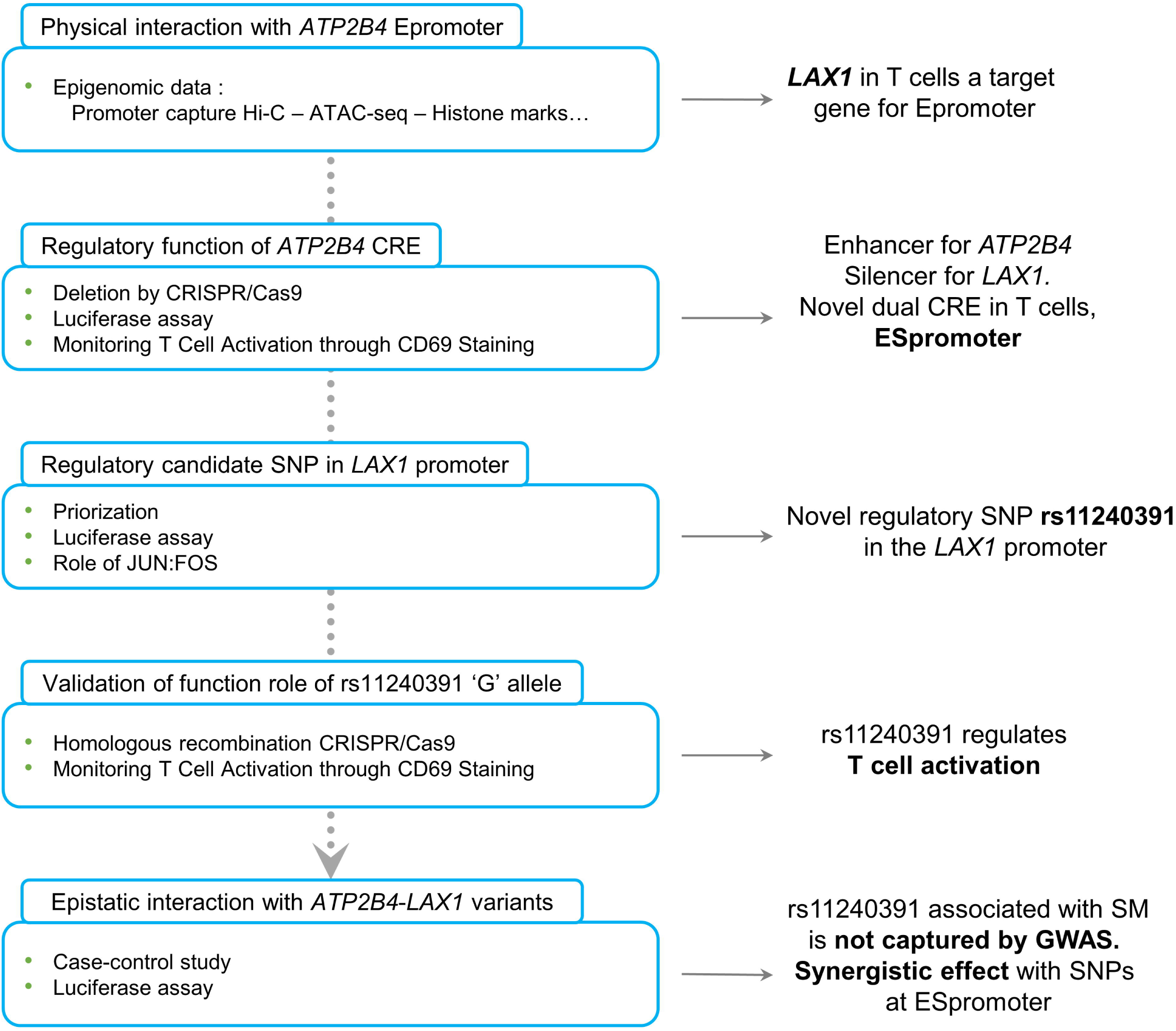
Workflow of bioinformatics and experimental analysis with the key findings, related to Figure 1-6.

**Figure S2.**
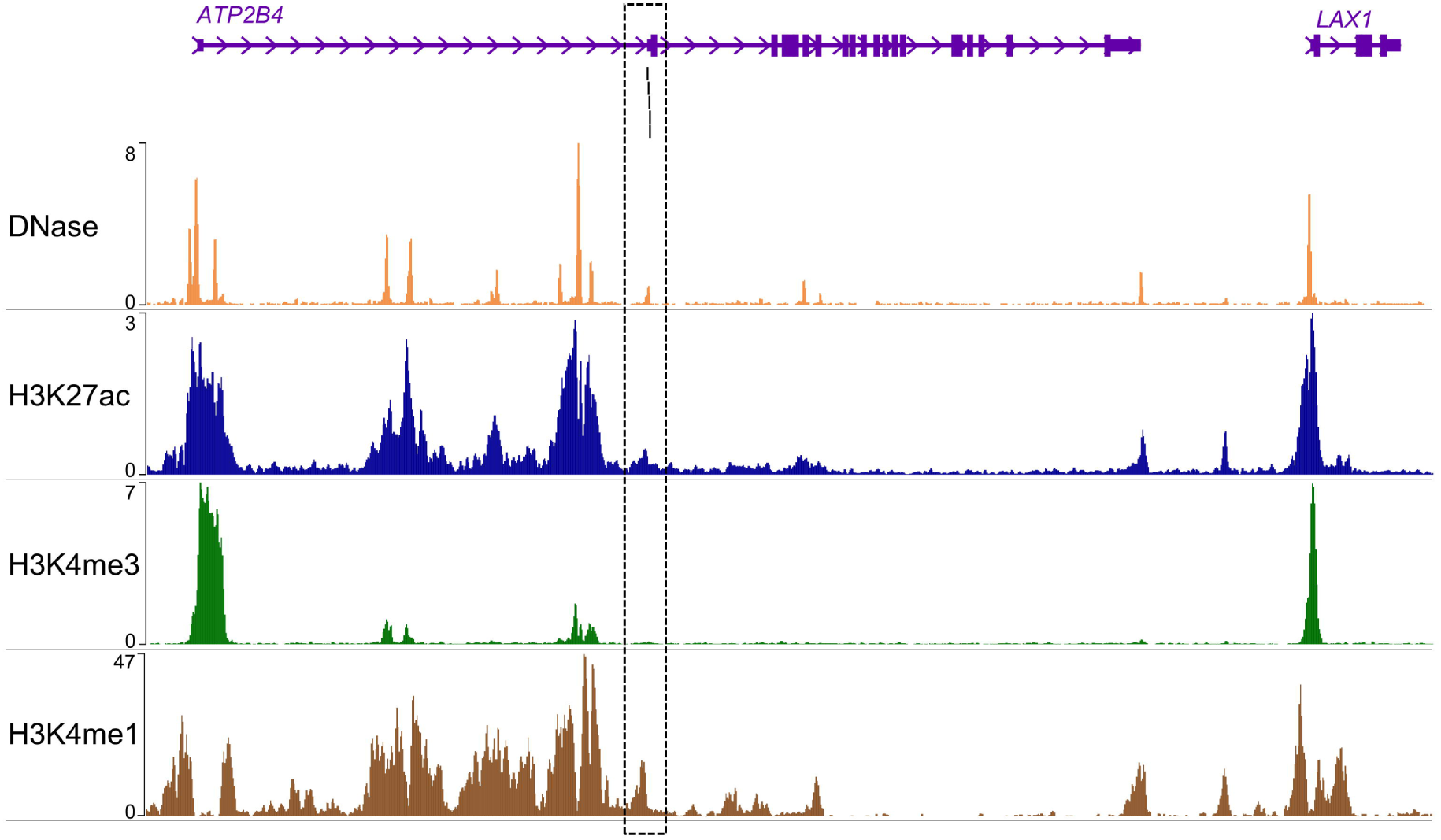
Epigenomic marks of Jurkat cell, related to Figure 1. WashU epigenome browser view of epigenomic data in CD4+ Naive Primary cells. The frame corresponds to the Epromoter. Black lines correspond to the 5 SNPs previously identified (rs11240734, rs1541252, rs1541253, rs1541254, and rs1541255).

**Figure S3:**
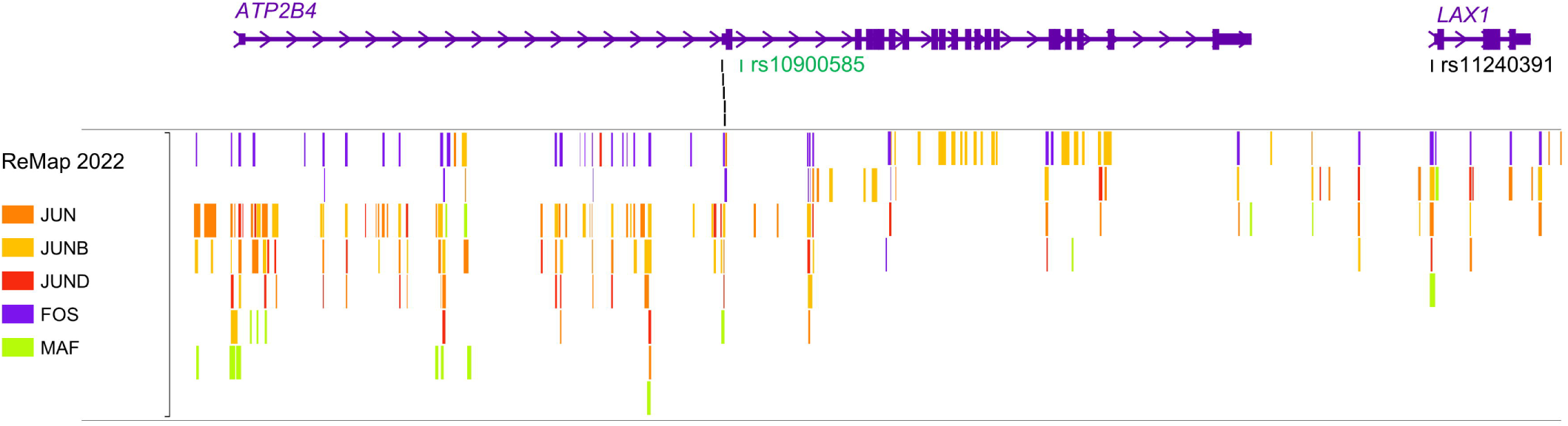
Identification of the FOS and JUN binding site on ESpromoter region and *ATP2B4* promoter, related to Figure 4. CHIP-seq peaks from ReMap2022 confirmed the binding of transcription factors, identified by RSAT, within the ESpromoter and *ATP2B4* promoters, in particular the FOS:JUN dimer corresponding to AP-1.

**Table S1: Description of the case-control cohort recruited in Senegal, related to Figure 6**

For each individual, the details indicated are identification number, phenotype (control: CTRL; Severe malaria: SM), age in years, sex (Female: F; Male: M), and the genotype for rs11240391.

**Table S2: Results of epistatic interaction between the 5 SNPs at ESpromoter and rs11240391, related to Figure 6**

**Table S3: Details of oligonucleotides used (name, source, identifier, and sequence), related to Key resources table**

## Notes

### Competing Interest Statement

The authors have declared no competing interest.

