## Supplemental table S2 for "Unraveling a novel dual-function regulatory element showing epistatic interaction with a variant that escapes genome-wide association studies"

Table S2: Results of epistatic interaction between the 5 SNPs at ESpromoter and rs11240391, related to Figure 6

| | Ctrl | SM | Odd ratio (95% CI) | P-value (two-tailed) |
|---|---|---|---|---|
| **Age** | NA | NA | 0.98 (0.97-0.99) | 0.002 |
| **Sex** | | | | |
| Male | 40 | 72 | 1 | - |
| Female | 38 | 38 | 0.57 (0.31-1.01) | 0.07 |
| ***LAX1*** | | | | |
| Risk genotypes (GG) | | | 1 | - |
| Protective genotype (TT+ TG) | 62 | 71 | 0.38 (0.19-0.75) | 0.005 |
| ***ATP2B4*** | | | | |
| Risk haplotype (Major/Major) | 26 | 64 | 1 | - |
| Protective haplotypes (Major/Minor + Minor/Minor) | 49 | 49 | 0.40 (0.22-0.74) | 0.004 |
| **Combination** | | | | |
| Risk genotypes *LAX1*/haplotype *ATP2B4* | 6 | 24 | 1 | - |
| Protective genotypes *LAX1*/haplotype *ATP2B4* | 39 | 28 | 0.18 (0.07-0.50) | 0.0008 |
| Interaction *LAX1* × *ATP2B4* | | | 0.33 (0.19-0.57) | 0.00006 |
