## Supplemental table S3 for "Unraveling a novel dual-function regulatory element showing epistatic interaction with a variant that escapes genome-wide association studies"

**Table S3: Details of oligonucleotides used (name, source, identifier, and sequence), related to Key resources table**

| Oligonucleotides | | |
|---|---|---|
| sgRNA | SEQUENCE | PAM |
| gRNA1 | TCCTCTACATTGGAGTTTAC | AGG |
| gRNA2 | TAGACTTCGGACGGCTACTC | GGG |
| gRNA3 | CCAATGTGCTAATGAAGCAC | AGG |
| ssODN sequence | SEQUENCE | |
| ssODN – G at rs11240391 | TATGTTTTCTTCTAGCAGATTAAGAGCTGA GCAGAGTTTCCTGTGCCCTGGGCTTCATTA GCACATTGGTGGTGTCGTTTCCGGTGACT GACTCTCTGTTT | |
| PCR primers | SEQUENCE | |
| F1 | GGCCACCCTTCAGATCACTT | |
| R1 | GCCTCCCTGTCTCAACTTCT | |
| F2 | TGAATCAGAAGAGGGTCCCG | |
| R2 | CGATCTCACCGGACATGGT | |
| F3 | CAAGAGTCTGGCCCGAGTTA | |
| R3 | TCTGCTGTTGAGATCGTCCA | |
| F4 | AGAAATTCTGAGAGCCCGGAG | |
| R4 | GATACCCACCGCGTACTCTG | |
| HPRT1 forward | GGGTGTTTATTCCTCATGGAC | |
| HPRT1 reverse | CTCCCATCTCCTTCATCACA | |
| Primers for mutagenesis | SEQUENCE | |
| Forward at rs11240391 | CTGTGCCCTGgGCTTCATTAG | |
| Reverse at rs11240391 | GAAACTCTGCTCAGCTCTTAATC | |
